# Target enrichment phylogenomics and biogeographic analyses unravel rapid radiation and reticulate evolution between Hainan-South China mainland -Vietnam in Section *Nintooa* (*Lonicera*, Caprifoliaceae)

**DOI:** 10.1101/2024.08.01.605766

**Authors:** Shi-You Zuo, Qing-Hui Sun, Diego F. Morales-Briones, Hong-Xin Wang, Jacob B. Landis, Hong-Yang Li, Hong-Jin Dong, Jun Wen, Hua-Feng Wang

## Abstract

The South China mainland and nearby islands are biodiversity hotspots. Section *Nintooa*, widely distributed across mainland China, Vietnam, and Hainan Island, exhibits a typical disjunct distribution pattern. However, the origins of the flora of Hainan Island and the phylogenetic relationships within Section *Nintooa* remain contentious. In this study, we collected 81 samples encompassing all recognized species of Section *Nintooa*, along with samples from other sections of *Lonicera*. We reconstructed the phylogenetic relationships using 491 orthologous nuclear genes and complete plastomes generated by target enrichment and genome skimming sequencing. Our nuclear gene-based species and concatenated trees support the monophyly of Section *Nintooa*. The species tree indicates that the Vietnamese and Hainan populations form sister clades. However, the plastome results indicate that Section *Nintooa* is polyphyletic, with subsection *Volubilis* forming a monophyletic group and subsection *Calcaratae* forming a sister clade with other members of subgenus *Chamaecerasus*. Our hybridization network analysis reveals extensive gene flow within Section *Nintooa*, whereas subsection *Calcaratae* shows no gene flow with subsection *Volubilis*, leaving the origin of *Calcaratae* unclear. *Lonicera* species from Hainan Island exhibit unstable phylogenetic positions and underwent a rapid radiation during the Miocene. Biogeographical results indicate that populations from Vietnam and Hainan both originated from mainland China. Overall, our findings enhance the understanding of the evolutionary diversification history of *Lonicera*.

## 1 Introduction

From the earliest studies of evolution, islands have provided a framework for understanding evolutionary processes (Wallace, 1869). Unlike oceanic islands, which generally lack connections to continents, fluctuations in sea level can repeatedly connect and isolate continental islands from the mainland. This process can promote the evolution of unique biota on continental islands (Chen et al., 2015), which may be highly susceptible to global climate change (Bellard et al., 2014; Crowl et al., 2015; Taylor and Kumar, 2016). Understanding the biogeographical history and local adaptations of species discontinuously distributed across continents and adjacent islands can provide insights into the evolution of continental island biota and guide conservation efforts under climate change.

Numerous continental islands can be found along the Pacific coast of southern China, the largest being Hainan Island. Hainan Island hosts approximately 3,800 species across 1,283 genera. The flora of Hainan Province is most similar to that of Guangxi Province in southern China and northern Vietnam (Zhu, 2016). The mountainous areas of the East Asian mainland and nearby continental islands are biodiversity hotspots (Myers et al., 2000; Tang et al., 2006), attracting significant research interest in recent years on the evolution of the region’s flora.

Phylogeographical studies on species distributions in mainland China, Japan, and the Korean Peninsula have elucidated the evolution of these temperate regions’ disjunct species (Chen et al., 2012; Lee et al., 2013; Qiu et al., 2011; Ye et al., 2017). However, the evolutionary and biogeographical history of the flora of south China mainland and its adjacent islands remains largely unexplored. One hypothesis suggests that the formation of Hainan Island and its flora is closely related to the southeastward drift of the South China Plate during the Cenozoic (Zhu, 2016, 2017), based on the similarity of the flora between southern China and the South China Plate. However, this hypothesis lacks consideration of the phylogeography of representative disjunct species. Pleistocene sea level fluctuations resulted in the recurrent formation of land bridges connecting the south China mainland and continental islands, potentially providing suitable habitats for species migration (Voris, 2000; Zhang et al., 2016)

Few biogeographical studies have explored the evolutionary history of disjunct species between the south China mainland and Hainan Island. Previous research has revealed various biogeographical patterns (Tian et al., 2018). For instance, Pleistocene climatic changes caused the differentiation and population expansion of the cotton bollworm (*Helicoverpa armigera*), linking populations in Taiwan and Hainan Island to those in southeastern China (Lin et al., 2014). Biogeographic patterns of *Engelhardia. roxburghiana* suggest that Hainan Island was in close biotic contact with Chiang-nan during the Eocene, spreading to Hainan Island from Chiang-nan via a land bridge during the late Eocene (Huang et al., 2024). These findings highlight the need for further investigation into the biotic assemblages on these continental islands along south China mainland.

The *Lonicera macrantha* complex, part of Section *Nintooa* of *Lonicera* (Caprifoliaceae), includes *Lonicera confusa*, *L. hypoglauca, L. macrantha, L. reticulata*, and *L. similis*. (Yang et al., 2011). Species of this complex are primarily distributed in Hainan Island, mainland south China, and northern Vietnam, representing a typical disjunct distribution between Hainan Island, mainland China, and northern Vietnam. This unique geographical distribution provides an ideal system for exploring species differentiation across the region. Studying the diversification history and dispersal routes of the *L. macrantha* complex in the mainland China-Hainan-Vietnam region can offer phylogenetic insights into the geographical origins of Hainan Island.

Section *Nintooa* of *Lonicera* comprises approximately 180 species distributed across northern Africa, Asia, Europe, and North America, with 57 species endemic to China (Rehder, 1903; Yang and Landrein, 2011). Section *Nintooa* includes twining vines with usually hollow stems. Small bracts and adjacent calyx tubes are mostly separate, rarely fused; corollas are bilabiate with often long tubes. Species of Section *Nintooa* have similar habitats and morphological characteristics, but can be distinguished morphologically. For example, *L. hypoglauca* can be easily distinguished by its mushroom-shaped glands (Figure 1 D3), while *L. reticulata* is readily identifiable due to the curling of its leaf margins (Figure 1 C3).

**Figure 1.**
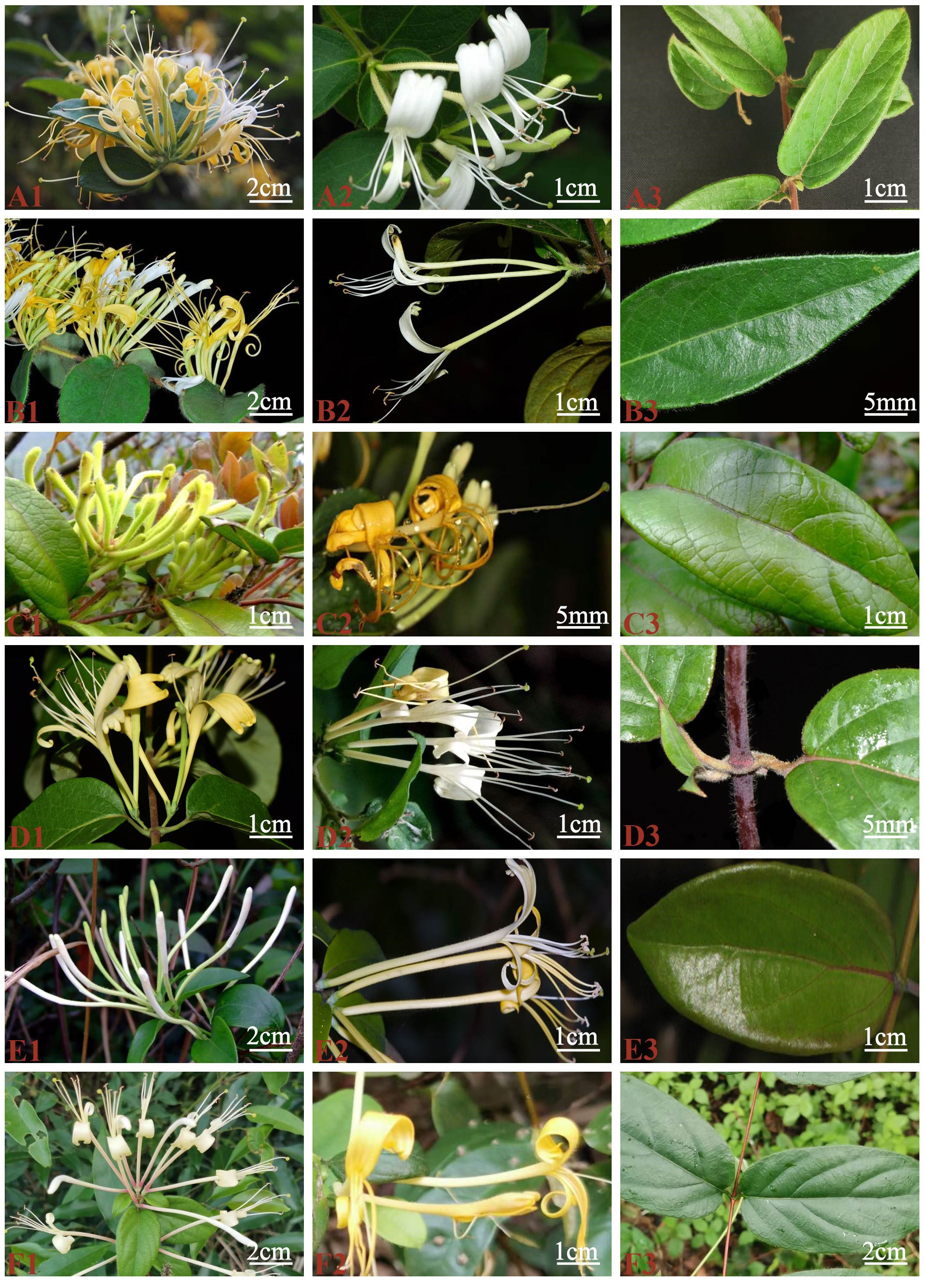
Morphology of Section Nintooa. A) Lonicera confusa, B) Lonicera macrantha, C) Lonicera reticulata, D) Lonicera longiflora, E) Lonicera calvescens

Since Linnaeus (1753) established *Lonicera*, the genus has presented numerous systematic challenges, particularly in infrageneric classification.Maximowicz, (1877) was the first to systematically classify Asian *Lonicera*, dividing the genus into three subgenera: *Caprifolium* (Tourn.) Maxim., *Chamaecerasus* (Tourn.) Maxim., and *Xylostron* (Tourn.). Rehder (1903, 1913) subsequently refined this classification by further subdividing the subgenus *Chamaecerasus* into four sections and 20 subsections. Nakai, (1938) developed a complex classification system for Japanese *Lonicera* species, involving both sections and subsections, identifying 15 sections and eight subsections. Later, Hara (1983) reclassified *Lonicera* into two subgenera: *Lonicera* and *Caprifolium* (Mill.) Dippel. Within subgenus *Chamaecerasus*, Hara established four sections: *Isika* (Anderson) Rehder, *Caeruleae* (Rehder) Nakai, *Lonicera*, and *Nintooa* (Sweet) Maxim. Hsu and Wang (1988) proposed a new classification for Chinese *Lonicera* species, dividing the genus into two subgenera. They recognized four sections and 12 subsections within the subgenus *Chamaecerasus* in China. In contrast, Yang et al. (2011) did not recognize taxa at the section level in their treatment of the Flora of China.

The phylogenetic relationships within the *L. macrantha* complex remain unresolved. Previous studies often had limited sampling and primarily utilized minimal chloroplast gene fragments or ITS and ETS sequences. Theis et al. (2008) inferred the relationships within *Lonicera* using ITS and sequences from five chloroplast non-coding regions (*rpoB-trnC, atpB-rbcL, trnS-trnG, petN-psbM, and psbM-trnD*), but their results did not support *Isika* and Section *Nintooa* as monophyletic groups. Dong and Peng (2014) investigated the interspecific relationships within the *L. macrantha* complex by combining phenotypic analysis with molecular evidence. They used two plastid DNA regions, psbA-trnH and trnS-trnG, in their phylogenetic analysis. However, their phylogenetic tree exhibited low support rates; only *L. macranthoides* was recognized, while the relationships among the other four species remained unresolved. Srivastav et al. (2023) utilized RADSeq to construct a well-supported phylogenetic tree, redefining the relationships within Section *Nintooa*. However, their study did not include *Lonicera calcarata*, leaving the phylogenetic position of Subsection *Calcaratae* unresolved. Yang et al. (2024) used chloroplast genomes and ITS and ETS sequences to establish a phylogenetic tree. Their results indicated that Section *Nintooa* was polyphyletic in the chloroplast tree, with Subsection *Calcaratae* nested within Section *Isika*. In contrast, in the ITS and ETS nuclear gene trees, Section *Nintooa* formed a monophyletic group, with Subsection *Calcaratae* being sister to Subsection *Volubilis*. This demonstrated a significant incongruence between the nuclear and chloroplast phylogenies. Overall, these studies highlight the need for further comprehensive research with increased sampling and the use of additional genetic markers to clarify the phylogenetic relationships within *Lonicera*. The inconsistencies and limitations observed in previous studies underscore the complexity of *Lonicera* phylogeny and the necessity for more robust methodologies.

The current study has two main objectives: First, to infer the origin of the *L. macrantha* complex on Hainan Island through biogeographical analyses. Second, to clarify the phylogenetic relationships within the *L. macrantha* complex by sampling populations of Section *Nintooa* in Hainan, South China mainland and Vietnam.

## 2 Materials and methods

### 2.1 Taxon sampling

We sampled 81 individuals from 30 species of Caprifoliaceae, including four sections (Section *Nintooa,* Section *Coeloxylosteum,* Section *Isika,* and Section *Isoxylosteum*). In addition, four genera from the Dipsacales (*Diabelia serrata*, *Diervilla sessilifolia*, *Dipelta floribunda*, *Weigela floribunda*) were used as outgroups. Species, vouchers, and GenBank accession numbers are provided in Table 1. We extracted DNA from dried leaf tissue using a modified cetyl-trimethylammonium bromide (CTAB) method (Doyle and Doyle, 1987) The quality and quantity of the DNA were assessed using an ultramicro spectrophotometer and 1% agarose gel electrophoresis. Each DNA sample was quantified with an Agilent 2100 BioAnalyzer. DNA samples with a minimum of 0.8 μg were selected for library preparation and whole-genome sequencing. Paired-end sequencing libraries with insert sizes of 300-500 bp were constructed and sequenced at the Beijing Genome Institute (BGI, Shenzhen, China) on the BGISEQ-500 platform. Raw reads were filtered and trimmed using SOAPfilter v2.2 with the following criteria: (i) read pairs were removed if the unknown base content in a single read exceeded 10% of its length or if low-quality bases (≤10) accounted for more than 40% of the read length and (ii) reads generated by PCR duplication were also filtered out.

**Table 1:**
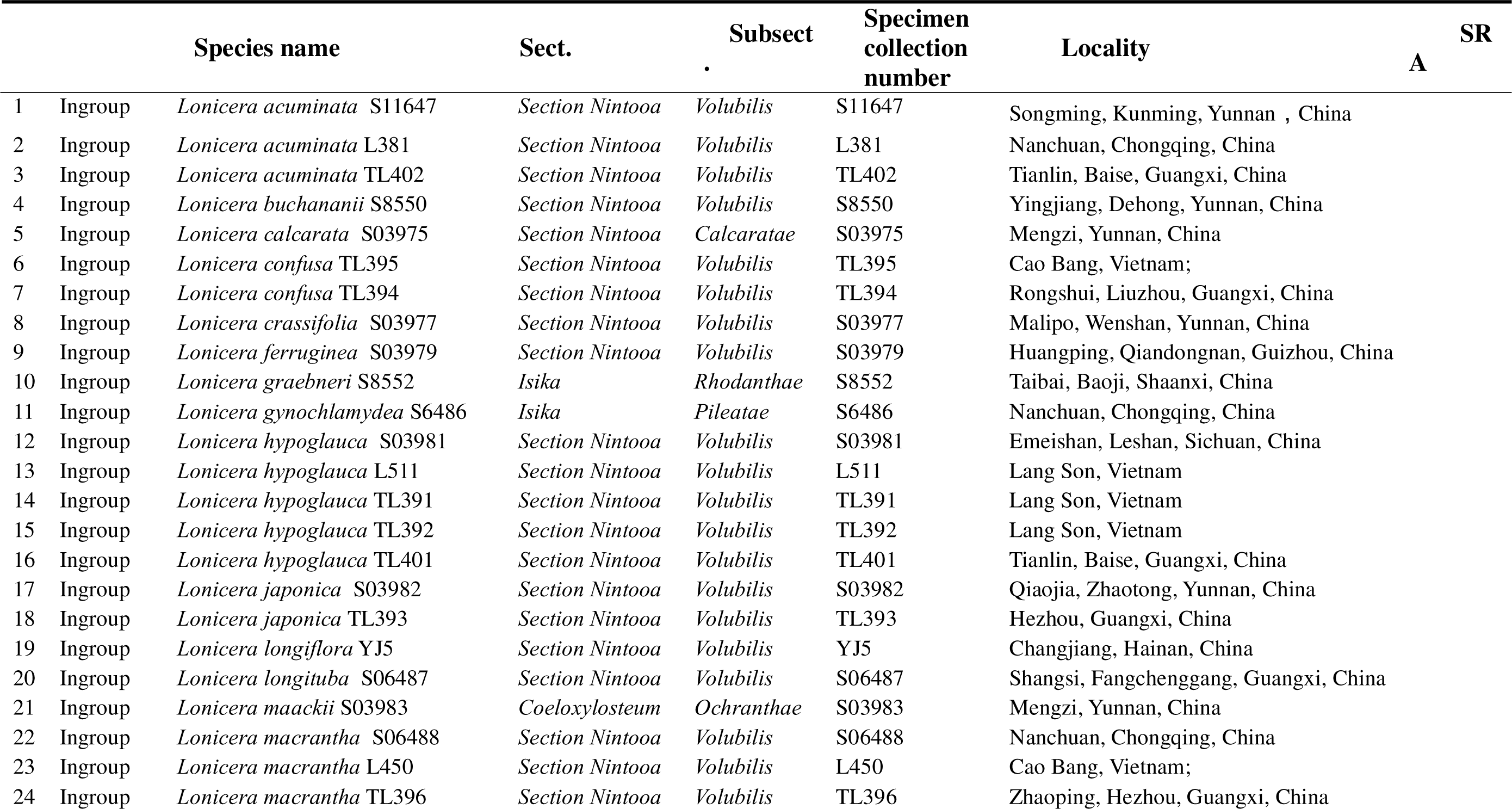
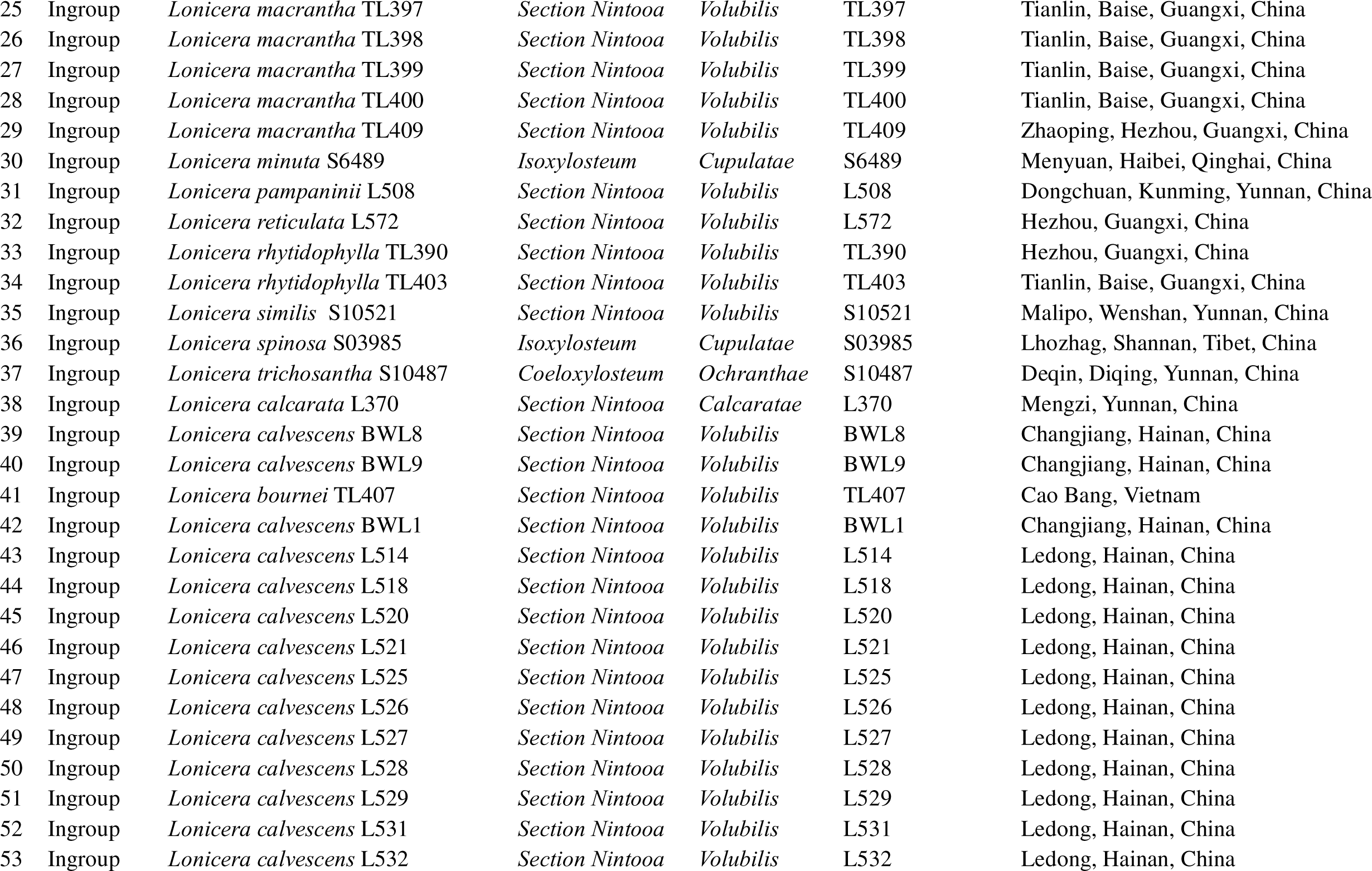
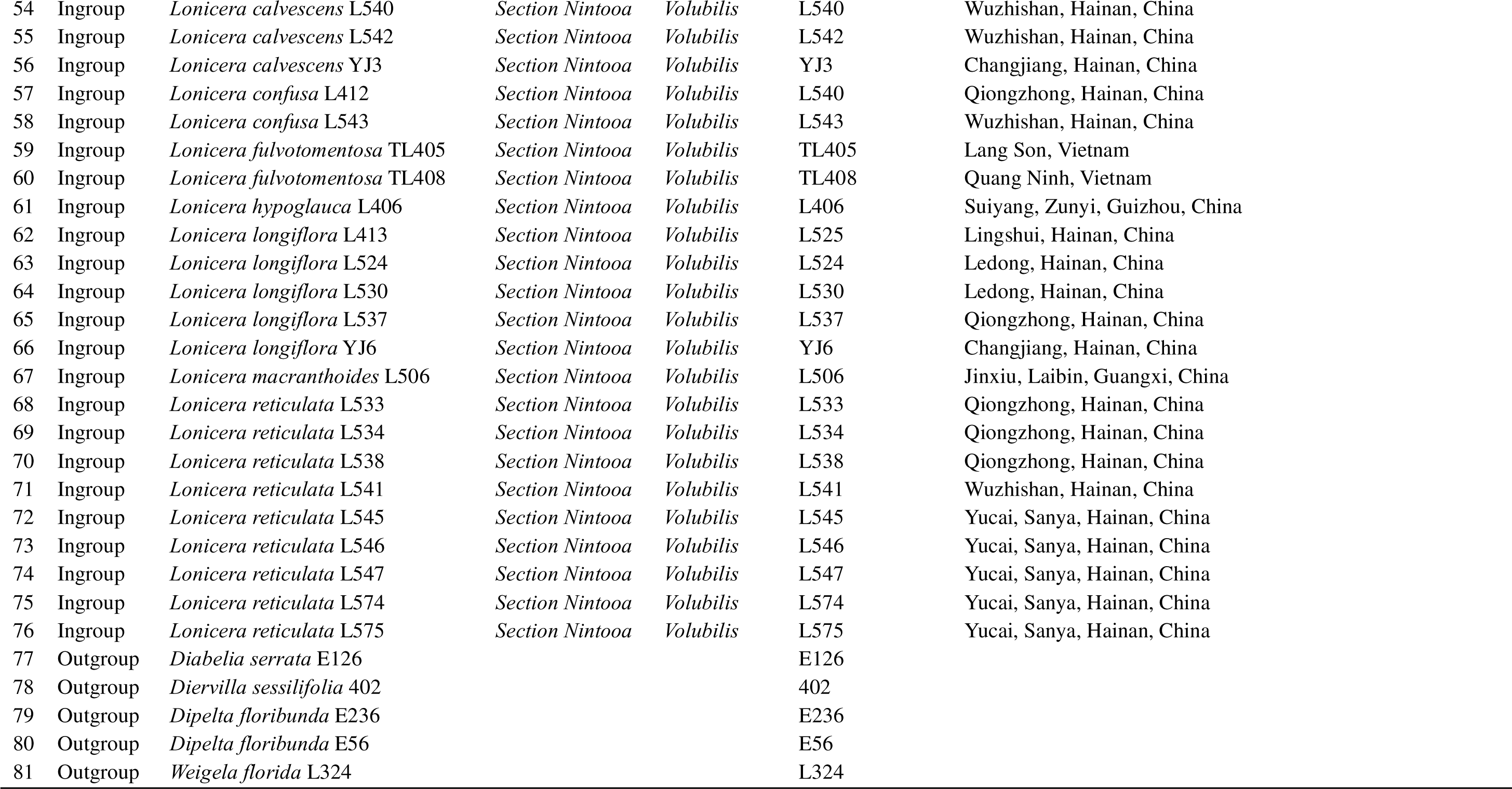
List of species and vouchers used in this study.

### 2.2 Read processing and assembly

We aimed to extract the 418 genes used in Caprifoliaceae s.l. (CAPRI418; Wang et al. 2021) and the Angiosperm 353 universal genes (A353;Baker et al., 2021)). We used CAPTUS v.0.9.91 (Ortiz et al., 2023) to trim the sequencing adaptors and low-quality bases, assemble the reads, and extract the nuclear loci using default parameters. For extraction of the CAPRI418 genes, we used as references the complete CDS from the original transcriptomes used by Wang et al. (2021) to develop the bait set. For the A353 genes, we used the extended set of references provided in CAPTUS. We extracted loci from both data sets using minimum percentages of identity and coverage of 75% and 50%, respectively. FASTA files were collected while retaining all paralog copies per sample. Finally, 50 genes that overlapped between the CAPRI418 and A353, were removed to keep only one copy, resulting in a combined 721-gene data set.

### 2.3 Orthology Inference

To infer orthologs for phylogenetic reconstruction, we followed a modified version of the methods described by Morales-Briones et al. (2022; https://bitbucket.org/dfmoralesb/target_enrichment_orthology). Initially, the untrimmed and unfiltered alignments from CAPTUS were re-aligned using MAFFT v7.520 (Katoh and Standley, 2013) with default parameters. Frameshifts were addressed by replacing “!” with gaps, and columns with more than 90% missing data were removed using Phyx (Brown et al., 2017).

Maximum likelihood (ML) homolog gene trees were inferred using RAxML v8.2.13 with the GTRGAMMA model and 100 bootstrap replicates. Monophyletic and paraphyletic tips belonging to the same taxon were masked, retaining tips with the most unambiguous characters in the trimmed loci alignments for each taxon, as described by Yang and Smith (2014). Spurious tips with unusually long branches were removed using TreeShrink v1.3.9 (Mai and Mirarab, 2018). TreeShrink was run twice with the quantile set to 0.01, using the output from the first run for the second to avoid over-trimming. FASTA files were generated from the output homolog trees, and the same steps as for the CAPTUS output alignments were followed: alignments were performed using MAFFT with default parameters, frameshifts were replaced with gaps, and columns with more than 90% missing data were removed using Phyx. To infer the final homolog gene trees, we used RAxML with the GTRGAMMA model, and assessed clade support with 1,000 ultrafast bootstrap (BS) replicates.

Orthology inference was carried out using the tree-based “monophyletic outgroup” (MO) method described by Yang and Smith (2014). The MO method searches for clusters with monophyletic ingroups rooted at the outgroups in the homolog trees, discarding those with duplicated taxa in the outgroups. Subsequently, it infers the orthologs from root to tip, retaining the ortholog subtree with the most taxa. For orthology inference, all *Lonicera* members were set as the ingroup, and the remaining taxa (*i.e.*, *Diabelia serrata*, *Dipelta floribunda*, *Weigela floribunda*, *Diervilla sessilifolia*) were set as the outgroup. Only ortholog groups with at least 21 ingroup taxa were retained. This resulted in a dataset of 81 samples and 491 ortholog trees.

We used GetOrganelle v1.7.5.1 (Jin et al., 2020) to assemble the chloroplast genomes from the clean reads of each species, employing default parameters (online manual https://github.com/Kinggerm/GetOrganelle). The target assembly graphs (GFA) were visualized using Bandage v0.8.1 (Wick et al., 2015) to assess the completeness of the final assembly. For each species, Before annotating the assemblies, we used Mauve v1.1.3 (Darling et al., 2004) to check the collinearity of the genomic sequences, selecting the sequences with consistent orientation for annotation. The chloroplast genome sequences were initially annotated with Geneious Prime v2023.0.1 (Kearse et al., 2012), using closely related species from GenBank as reference sequences. Further manual editing was conducted to refine the start codons, stop codons, and intron/exon boundaries.

### 2.4 Phylogenetic analyses

#### 2.4.1 Nuclear dataset

We used concatenation and coalescentLbased methods to infer the phylogeny of Section *Nintooa*. A concatenated alignment was produced using the clean ortholog alignments (following the Phyx step), retaining 491 orthologs. We estimated a ML tree with RAxML v.8.2.13 (Stamatakis, 2014). Clade support was assessed with 1,000 rapid bootstrap replicates (BS). For the coalescent-based approach, we first inferred ML gene trees for the 491 individual orthologs with RAxML using the GTRGAMMA model. We then estimated a species tree with ASTRALLIII v.5.7.1 (Zhang et al., 2018). Local posterior probabilities (LPP; (Sayyari and Mirarab, 2016) were used to assess clade support. We used Quartet Sampling (QS) (Pease et al., 2018) to distinguish strong conflict from weakly supported branches.

#### 2.4.2 Plastome dataset

We conducted Maximum Likelihood (ML) analysis using RAxML. Initially, the plastome sequences were aligned using MAFFT v.7.407 (Katoh and Standley, 2013), and columns with more than 90% missing data were removed and using the GTR+I+G model with 1000 bootstrap replicates. Additionally, we used Quartet Sampling (QS) with 1000 replicates to further evaluate the reliability of the phylogenetic tree.

### 2.5 Species network analysis

Since our network analysis primarily focused on potential network structures among the major clades of Section *Nintooa*, we reduced the 81-sample data set to one outgroup and 14 ingroup taxa representing all major clades based on nuclear analyses. Additionally, to investigate the causes of the phylogenetic position inconsistency of subsection *Calcaratae*, we selected seven species (each representing different sections) for network analysis. Given the computational restrictions of computing phylogenetic networks in PhyloNet we used the InferNetwork_MPL command(Yu and Nakhleh, 2015) based on the maximum pseudo-likelihood (MPL) measure. For all tests, the number of optimal output networks was set to 10. The “po” option was specified to optimize branch lengths and inheritance probabilities of the inferred networks under full likelihood. The network with the smallest pseudo-likelihood value was chosen as the best network. We visualized the optimal network using Dendroscope v3.8.8 (Huson and Scornavacca, 2012).

### 2.6 Divergence time estimation

The nuclear dataset was used to estimate the divergence time of *Lonicera* with BEAST v.2.4.0 (Bouckaert et al., 2014). We employed the GTR+G substitution model and a Birth-Death model tree prior. Other parameters were left at default settings. The root age was set to 78.9 Ma (mean 78.9 Ma, normal prior distribution 76.3–82.2 Ma) following Li et al. (2019). We selected two fossils as calibration points. First, we selected the fossil seeds of *Weigela* Thunb. from the Miocene and Pliocene in Poland (LańcuckaLRodoniowa, 1967) and the Miocene in Denmark (Friis, 1985) to constrain the stem age (offset 23.0 Ma, lognormal prior distribution 23.0–28.4 Ma). Second, we set a lognormal prior for the divergence at the stem of *Diabelia* and its sister clade *Dipelta* (offset 36 Ma, log-normal prior distribution 34.07–37.20 Ma).

We performed multiple MCMC runs for a total of 7 billion generations sampling every 1000 generations. Stationarity was checked with Tracer v.1.7 (Rambaut et al., 2018). All ESS values exceeded 200, and the first 25% of trees were discarded as burn-in with LogCombiner from multiple runs, and then determined the maximum clade credibility (MCC) tree with mean heights for the nodes in TreeAnnotator. The 95% highest posterior density (HPD) intervals were viewed in FigTree v.1.4.4 (Drummond et al., 2012).

### 2.7 Ancestral area reconstructions

To infer the ancestral distribution of species, we employed the R package BioGeoBEARS v.0.2.1, along with a time-calibrated phylogeny (Matzke, 2013). The ancestral area reconstruction was divided into three main geographic regions: (A) Hainan, (B) Mainland China, and (C) Vietnam. BioGeoBEARS reconstructs ancestral geographic distributions on phylogenies while testing for the best-fit range evolution model. It incorporates the basic assumptions of three widely used models in historical biogeography: DEC (Dispersal-Extinction-Cladogenesis; Ree and Smith, 2008), DIVA (Dispersal-Vicariance Analysis; Ronquist, 1997), and BayArea (Bayesian Inference of Historical Biogeography for Discrete Areas; Landis et al., 2013), allowing for direct comparison within a Maximum Likelihood framework. These models encompass a range of biogeographic processes such as vicariance, sympatric speciation, range expansion, and contraction. They can also be combined with a founder-event (”jump”) speciation model specified by the parameter “j” (Matzke, 2014).

We conducted multiple BioGeoBEARS runs using six models: DEC, DEC+J, DIVALIKE, DIVALIKE+J, BAYAREALIKE, and BAYAREALIKE+J, to identify the best-fitting model for our data. The input tree was a time-calibrated maximum clade credibility tree obtained from our divergence time estimation analysis. Each model was evaluated using Akaike Information Criterion (AIC) and ΔAIC values, with the model having the lowest AIC considered the best fit for our data.

Additionally, we utilized the Bayesian binary Markov chain Monte Carlo (BBM) method for reconstructing ancestral states using the program RASP (version 4.2; Yu et al., 2015). This method was used to reconstruct ancestral states over 5000 generations with nine hot chains and one cold chain. Each sample was assigned to its respective region based on its contemporary distribution range.

## 3. RESULTS

### 3.1 Phylogenetic results

Comparing the results of the nuclear and chloroplast data sets, although the topologies received good support, the positions of branches formed by *Lonicera* from Vietnam and mainland China were found to be unstable across different phylogenetic analyses (Figure 3).

The species tree shows a topology similar to the nuclear concatenated tree (Figure 2, Figure S1). Section *Isika*, Section *Coeloxylosteum*, and Section *Isoxylosteum* form a clade, which are sister to Section *Nintooa* (LPP=1) with strong support with QS (QS=1/NA/1). Furthermore, we demonstrate strong support (LPP=1, QS=0.7/1/0.98) for the sister relationship between the subsection *Calcaratae* and subsection *Volubilis*. By considering the geographical origins of the samples, we found that populations of *Lonicera* from mainland China formed polytomies. We found that *Lonicera* species from Dacunshan, Guangxi formed a polytomy; where *L. acuminata* TL402, *L. macrantha* TL397, *L. macrantha* TL400, and L. rhytidophylla TL403 clustered together, sister to a clade comprising *Lonicera* species from Hainan Island and Vietnam (LPP=0.76, QS=-0.24/0.16/0.88). However, *L. macrantha* TL398 and *L. macrantha* TL399 from Dacunshan, Guangxi, clustered with *L. acuminata* L381 and *L. pampaninii* L508 from Yunnan, forming a clade (LPP=0.68) sister to a clade comprising *Lonicera* populations from Yunnan and Chongqing, strongly supported by local posterior probabilities (LPP=1). Additionally, specimens from eastern Guangxi (*L. confusa* TL394, *L. japonica* TL393, *L. macrantha* TL396, *L. macrantha* TL409) and eastern Guizhou (*L. ferruginea* S03979) formed a clade, strongly supported by local posterior probabilities(LPP=1). We also found that *Lonicera* species from Hainan Island formed a distinct clade, sister to *L. fulvotomentosa* TL405 from Vietnam, strongly supported by local posterior probabilities (LPP=0.91) and moderate QS support (QS=0.24/0.88/0.97). However, within Hainan Island, the phylogenetic relationships of *Lonicera* are complex, with species from Apart from *L. fulvotomentosa* TL405 clustering with Hainan Island *Lonicera*, other species from Vietnam grouped together with moderate local posterior probability support.

**Figure 2.**
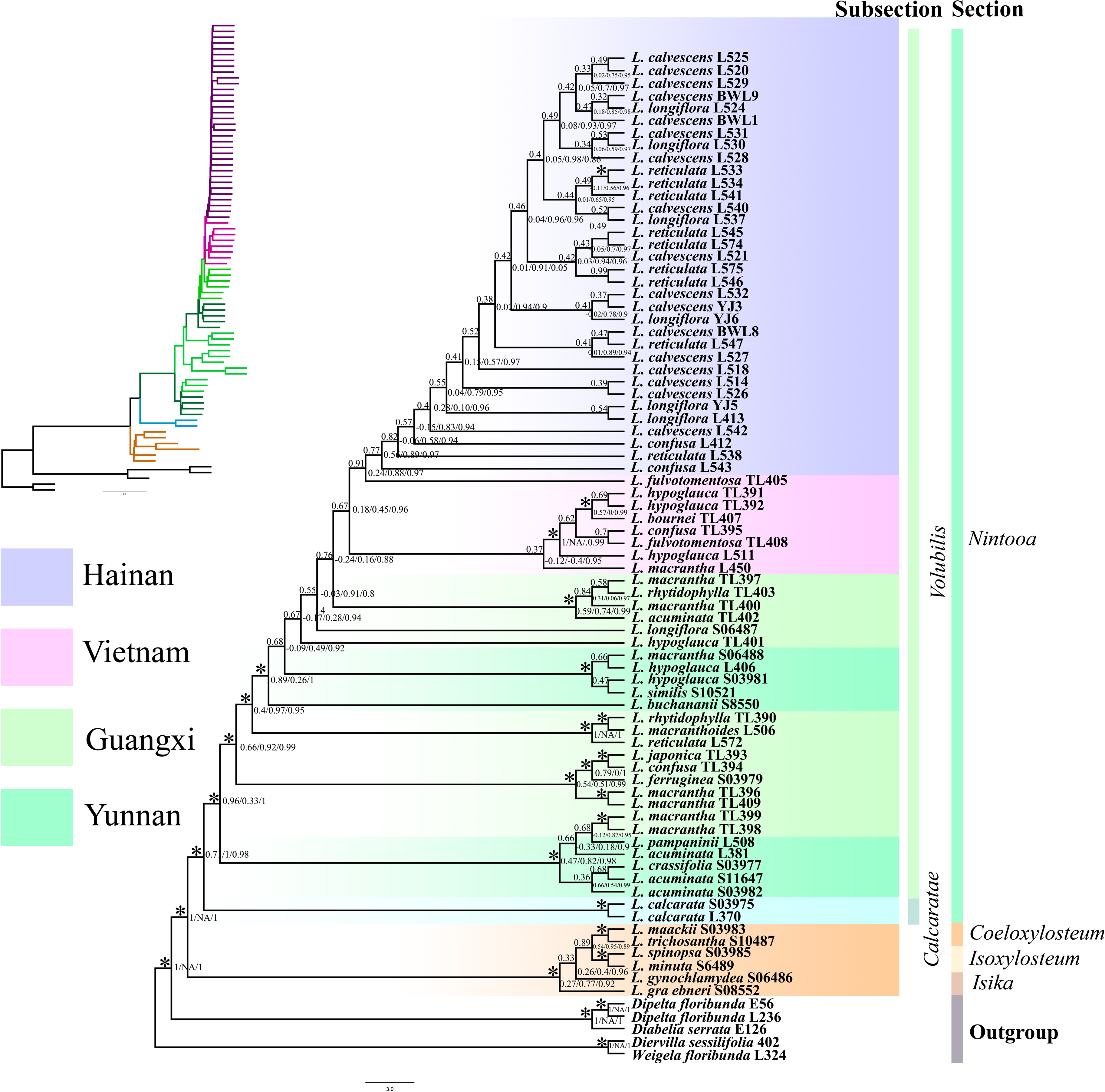
The nuclear species tree with ASTRAL-III. Clade support is depicted as: Quartet Concordance (QC)/Quartet Differential (QD)/Quartet Informativeness (QI)/Local posterior probabilities (LPP).

The concatenated nuclear gene tree (Figure 3, Figure S2) exhibits a topology similar to that of the species tree, with strong or full support (BS>85) for the major phylogenetic relationships. Within the concatenated nuclear gene tree, we find Section *Nintooa* recovered as a monophyletic group, with maximum support and complete QS backing (QS =1/NA/1). Within Section *Nintooa*, we observe the *Volubilis* and *Calcaratae* subclades forming sister clades, receiving full bootstrap support (BS=100) and strong QS backing, along with signals for potential alternative topologies (QS =0.66/0.98/0.98).

**Figure 3.**
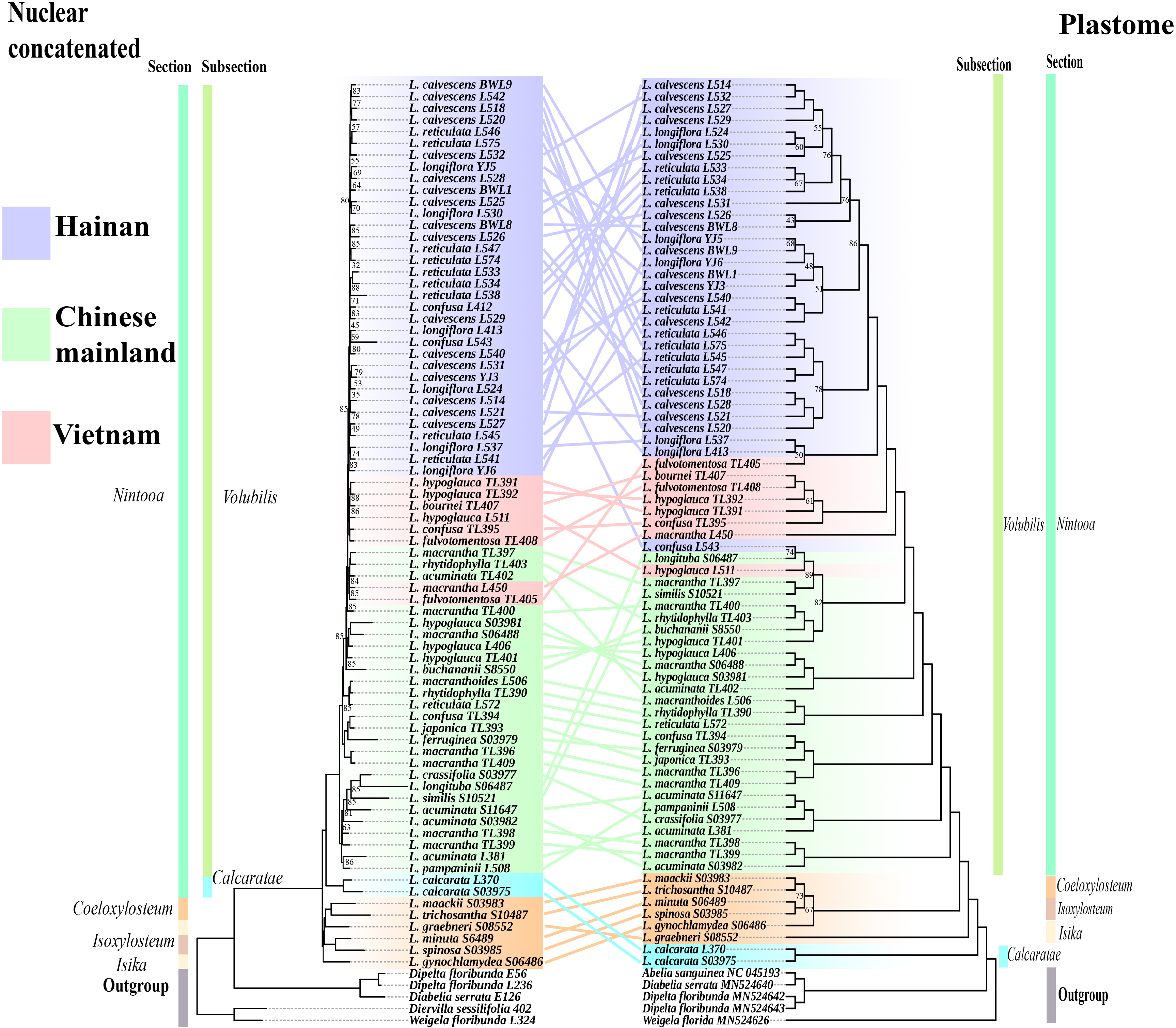
Tanglegram of the nuclear concatenated (left) and plastid (right) phylogenies. Dotted lines connect taxa between the two phylogenies. Support values are shown above branches. Values with bootstrap (BS) ≥ 90 were not displayed. Major taxonomic groups or main clades in the family as currently recognized are indicated by branch colors as a visual reference to relationships.

Considering the geographical origins within the *Volubilis* subclade, although all *Lonicera* samples from Hainan Island cluster into a major branch, the Hainan Island *Lonicera* lack strong support, with generally low BS and QS support rates. Only a few *Lonicera* samples received robust support; for example, *L. calvescens* L528 from Jianfengling (Ledong) was nested within *L. calvescens* BWL1 and *L. longiflora* YJ5from Bawangling (Changjiang), albeit with relatively lower support (BS=64). This clade then forms a sister clade with *L. calvescens* L525 and *L. longiflora* L530 from Jianfengling, receiving strong support (BS=93) but weaker QS support and clear signals for alternative topologies (QS=0.088/0.85/0.97).

*Lonicera* species from Vietnam form two polytomies: *L. bournei* TL407; *L. hypoglauca* TL391; *L. hypoglauca* TL392; forming a clade with *L. confusa* TL395 and *L. hypoglauca* L511, strongly supported with robust QS support and potential signals for alternative topologies (QS=0.83/0.14/0.99). Two other samples from Vietnam, *L. fulvotomentosa* TL405 and *L. macrantha* L450, form a clade (BS=85,QS=-0.32/0.045/0.99), which sister to a clade comprising three species from Guangxi, *L. acuminata* TL402, *L. macrantha* TL397, and *L. rhytidophylla* TL403, receiving strong support (BS=84) but facing QS opposition and clear signals for alternative topologies (QS=-0.16/0.36/0.95).

Branching patterns of *Lonicera* from mainland China do not strongly correlate with geography with species from different regions often forming clusters with robust support. However, some *Lonicera* from Guangxi form a well-supported clade, with *Lonicera confusa* TL394, *Lonicera japonica* TL393, and *Lonicera ferruginea* S0397 forming a clade with strong support (BS=100), which then forms a sister clade with *L. macrantha* TL396 and *L. macrantha* TL409, receiving full support (BS=100) but only moderate QS support and no signals for alternative topologies (QS=0.55/0.57/1).

Based on the plastome tree *Lonicera* form a polytomy (Figure 3 and Figure S3). We found that the *Calcaratae* subclade is sister to the *Isika*, *Coeloxylosteum*, *Isoxylosteum*, and *Volubilis* subclades, with maximum support (BS=100, QS=1/NA/1). *Lonicera* species from the *Volubilis* subclade are sister to the *Isika, Coeloxylosteum*, and *Isoxylosteum* subclades, with full support (BS=100, QS=1/NA/1). Furthermore, *L. fulvotomentosa* TL405 from Vietnam and those from Hainan Island form a sister relationship, albeit with low support (QC<0.5, BS<50). Apart from *L. fulvotomentosa* TL405; *L. hypoglauca* L511 and *L. macrantha* L450, the remaining *Lonicera* samples from Vietnam form a clade, sister to *Lonicera* from Hainan Island, with strong support (BS=100, QS=1/0/0.98). *L. confusa* L543 and *L. longituba* S06487 from Mainland China form a clade with weak support (BS=74), which is sister to *L. hypoglauca* L511 from Vietnam, with strong BS support (BS=100) and relatively reliable QS support (QS=0.89/0/0.96).

### 3.2 Divergence time estimation

The common ancestor of *Lonicera* dates to the middle Eocene at 42.55 million years ago (Ma) (95% HPD = 31.44-52.11 Ma)(Figure 5). Subsequent divergences led to the differentiation of *Volubilis* and *Calcaratae* around 37.04 Ma (95% HPD = 31.44-52.11 Ma). Combining inferred divergence times with geographical data, *L. confusa* TL394, *L. japonica* TL393, *L. ferruginea* S03979, *L. macrantha* TL396, and *L. macrantha* TL409 from Guangxi Province diverged 19.75 Ma. The common ancestor of *Lonicera* from Hainan Island, Vietnam, and parts of Guangxi Province date to the early and middle Miocene at 15.74 Ma (95% HPD = 11.58-19.28 Ma), with the divergence of *Lonicera* from Hainan Island dating to 14.04 Ma (95% HPD = 10.62-17.04 Ma). Meanwhile, the divergence time for *Lonicera* species from Vietnam is estimated to be 14.15 Ma (95% HPD = 9.89-17.69 Ma).

### 3.3 Species network analysis

In the hybrid network analysis of the 14 samples (Figure 5A), we identified an optimal hybrid network involving five reticulation events. The first event involved *Lonicera* species from Vietnam, where *L. hypoglauca* L511 is hypothesized to have experienced gene flow with *L. bournei* TL407 and *L. macrantha* L450, contributing 31.7% and 68.3%, respectively, to their genetic makeup. The second hybridization event involved *L. calvescens* L525 (Ledong, Hainan Island), with a major genetic contribution from *L. calvescens* L540 (Wuzhishan, Hainan Island) at 80.3%. The third reticulation event involved *L. confusa* L543 (Wuzhishan, Hainan Island), with gene flow from *L. calvescens* L525 at 90.1%, and a minor contribution from the branch comprising *L. confusa* TL394 and *L. macrantha* TL409 at 0.99%. The fourth hybrid network concerned *Lonicera* from Vietnam and Hainan Island, primarily inheriting genetic material from their ancestors (65.2%). The fifth hybridization event involved *Lonicera* species from Vietnam, Hainan Island, and two mainland Chinese regions, revealing a predominant genetic contribution from *Lonicera* species in Yunnan, China (50.8%), with the remainder inherited from their ancestors (49.2%). To understand why the phylogenetic positions of *L. calcaratae* differ markedly between the nuclear and chloroplast trees, we selected different groups as input for hybrid network analysis. The results, shown in Figure 5B, indicate that the formation of *L. calcaratae* is inferred to be the result of reticulate events involving multiple gene flow and hybridization events. The primary genetic contribution comes from *L. macrantha* (90%).

**Figure 4.**
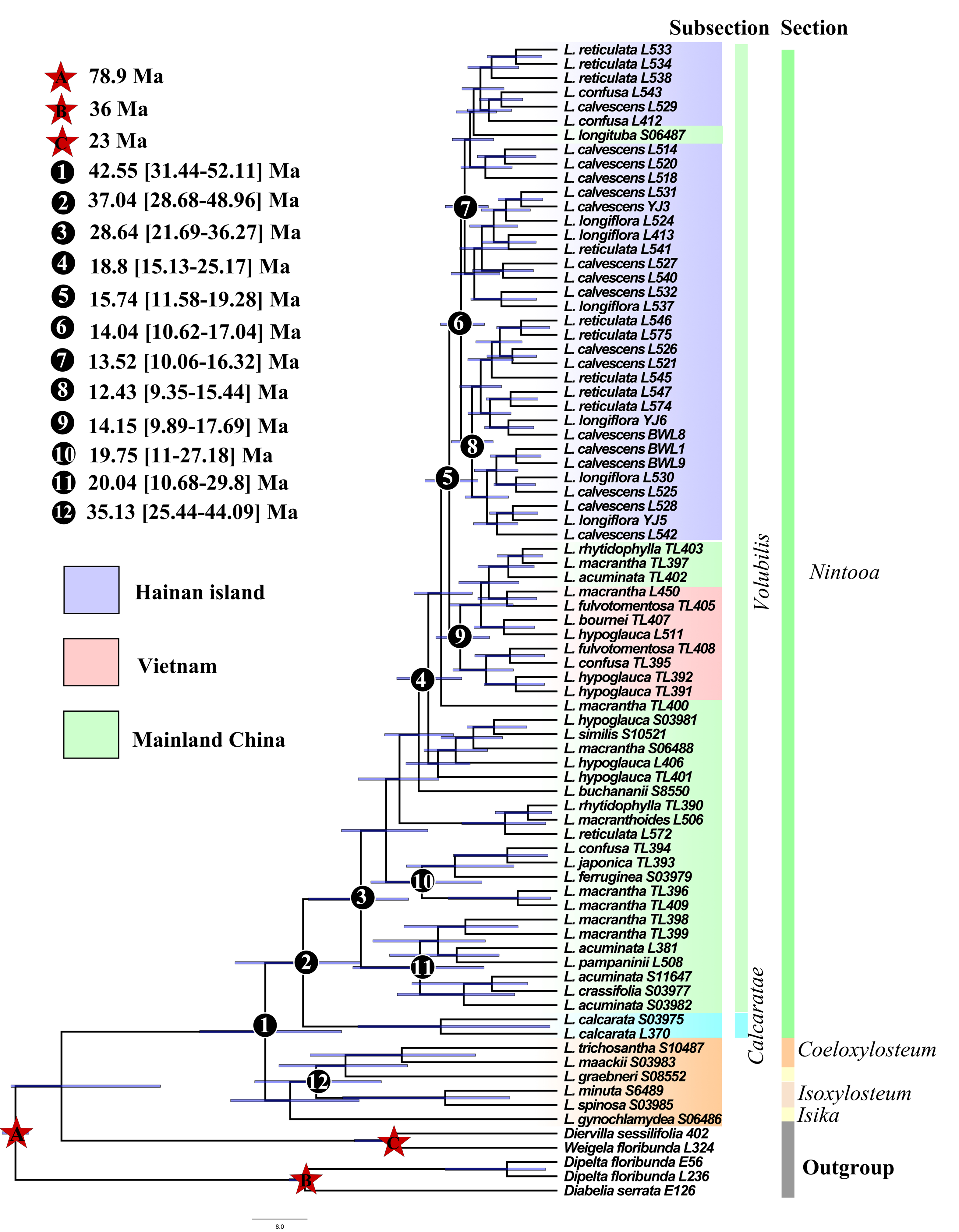
BEAST analysis of divergence times based on nuclear data. Calibration points are indicated by stars. Numbers 1–9 represent major divergence events in *Lonicera*. Mean divergence times and 95% highest posterior densities are provided for each major clade.

**Figure 5.**
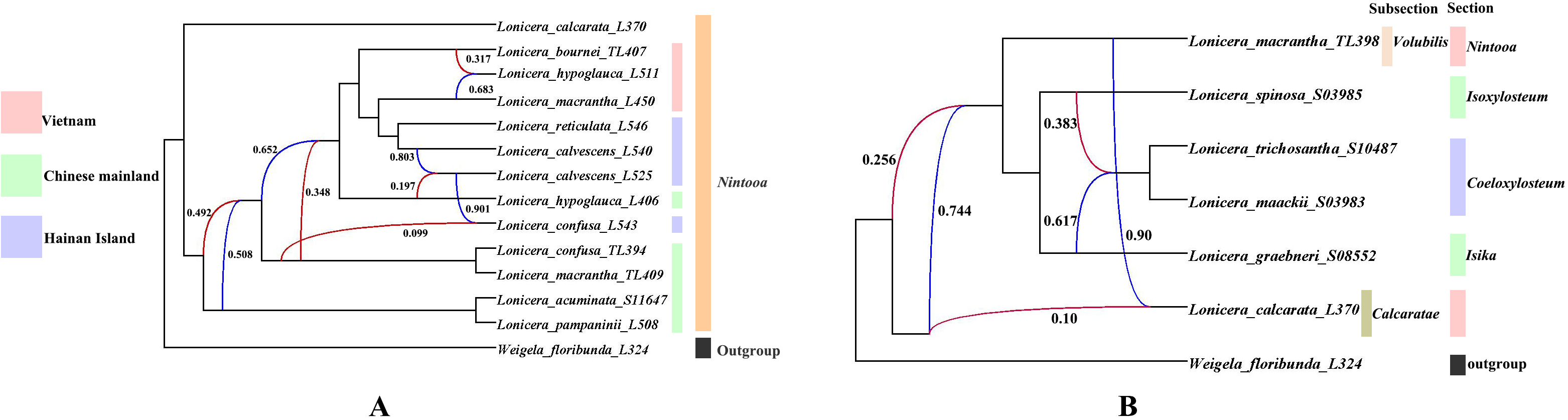
Best supported species networks inferred with PhyloNet for the (A) 14Ltaxa, and (B) 7Ltaxa. Numbers next to the inferred hybrid branches indicate inheritance probabilities. Blue curved lines represent major hybrid edges. Red curved lines represent minor hybrid edges (edges with an inheritance contribution <0.50).

### 3.4 Ancestral area reconstruction

In addressing the origin of the *Lonicera* group, we employed two analytical methods, BioGeoBEARS and RASP. Within BioGeoBEARS (Figure 6, Figure S4), our analyses indicated that the DEC+J model provided the best fit, with the lowest AIC value of 46.14013 (Table 4). The jump parameter (J) significantly improved the model fit, suggesting that both gradual dispersal and long-distance dispersal events (founder-event speciation) played crucial roles in the biogeographic history of *Lonicera* species (Table 4).

**Figure 6.**
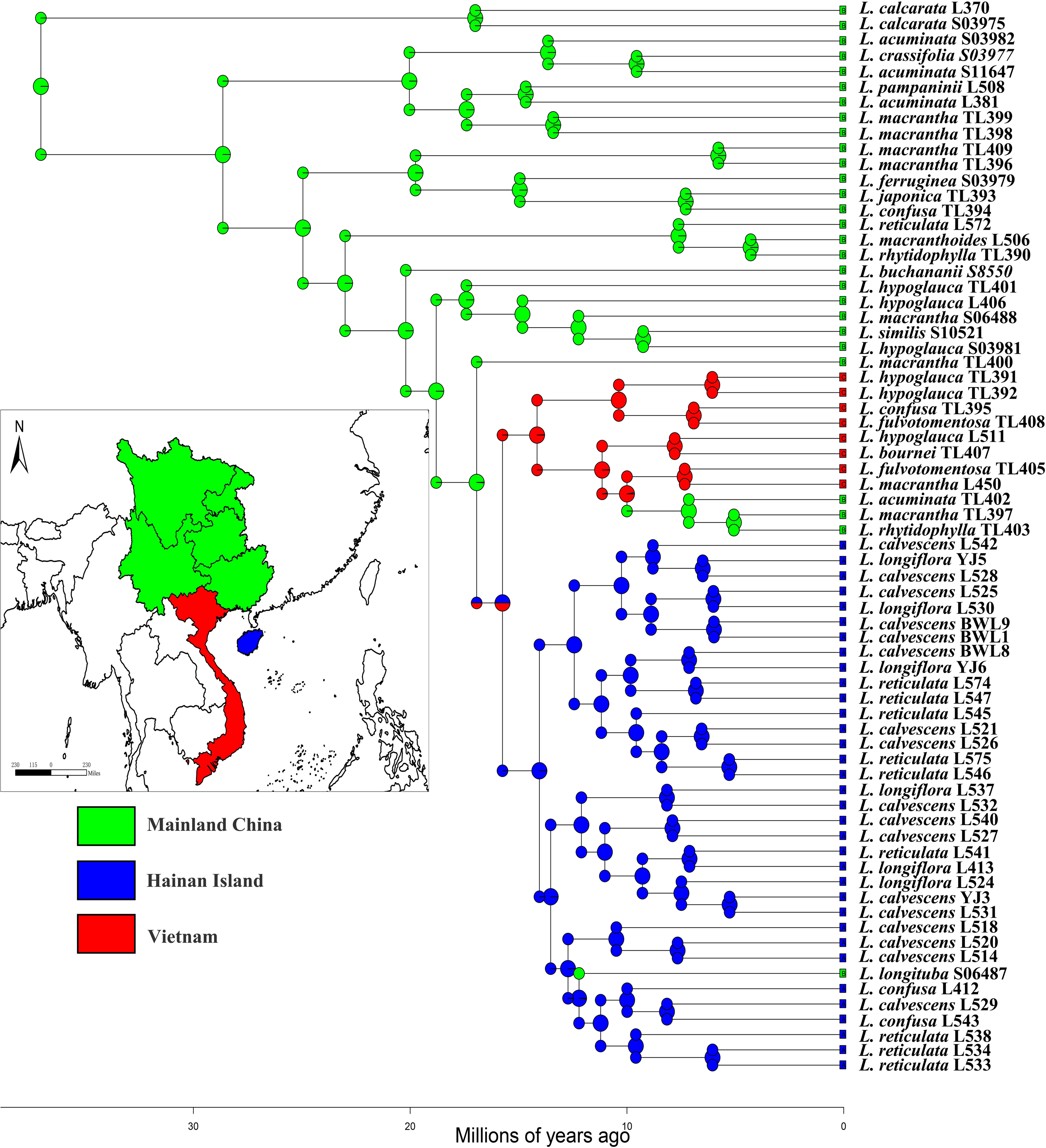
The DEC+J model in BioGeoBEARS to infer the ancestral range of Section *Nintooa*. The geographic distribution areas are delineated as follows: (A) Hainan Island; (B) Mainland China; (C) Vietnam.

**Table 2.**
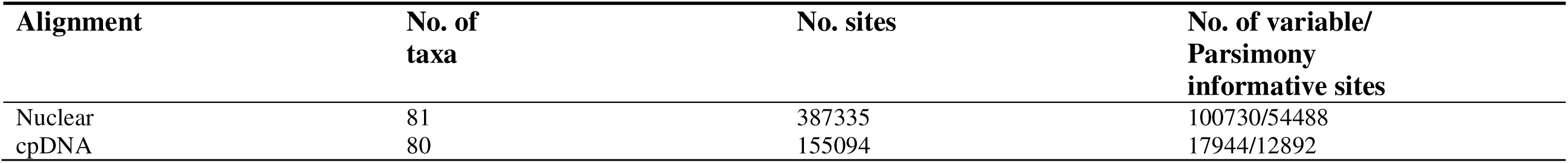
Dataset statistics, including the number of taxa, number of characters, number of PI characters.

**Table 3.**
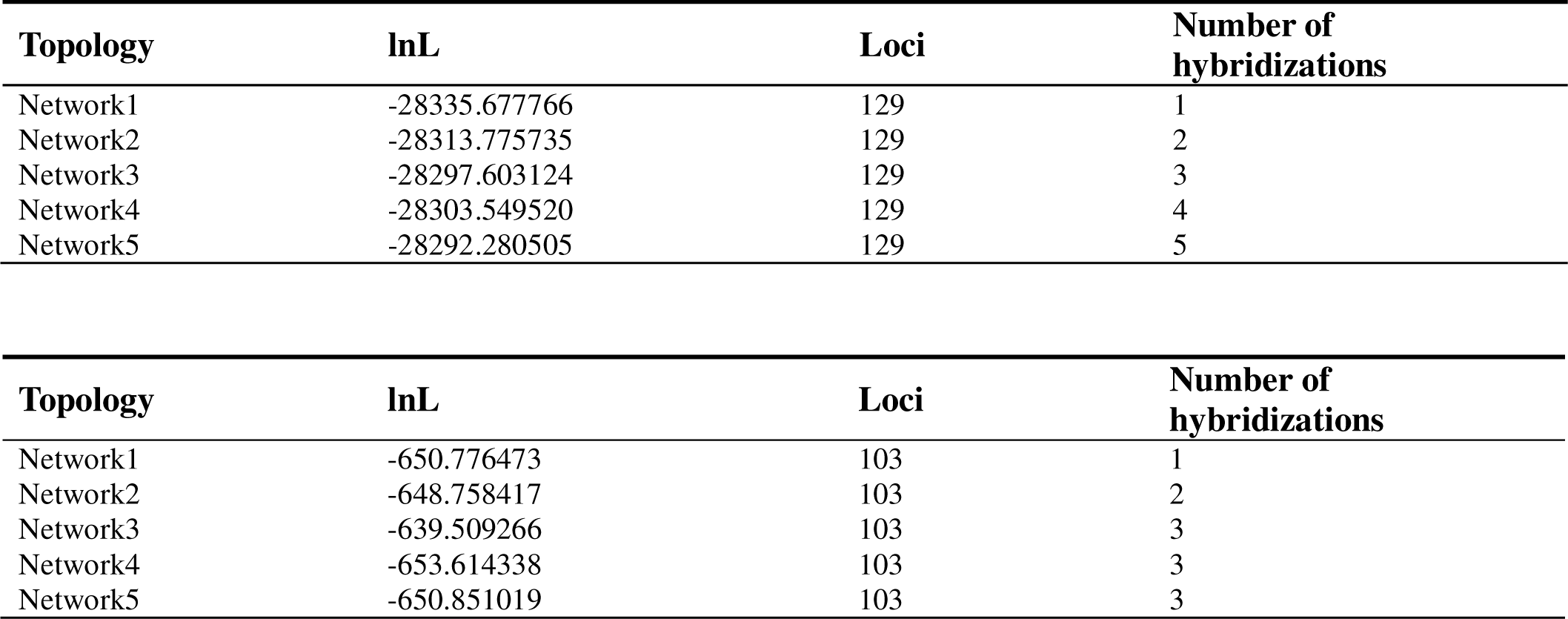
Model selection of the different species networks and bifurcating trees.

**Table 4.**
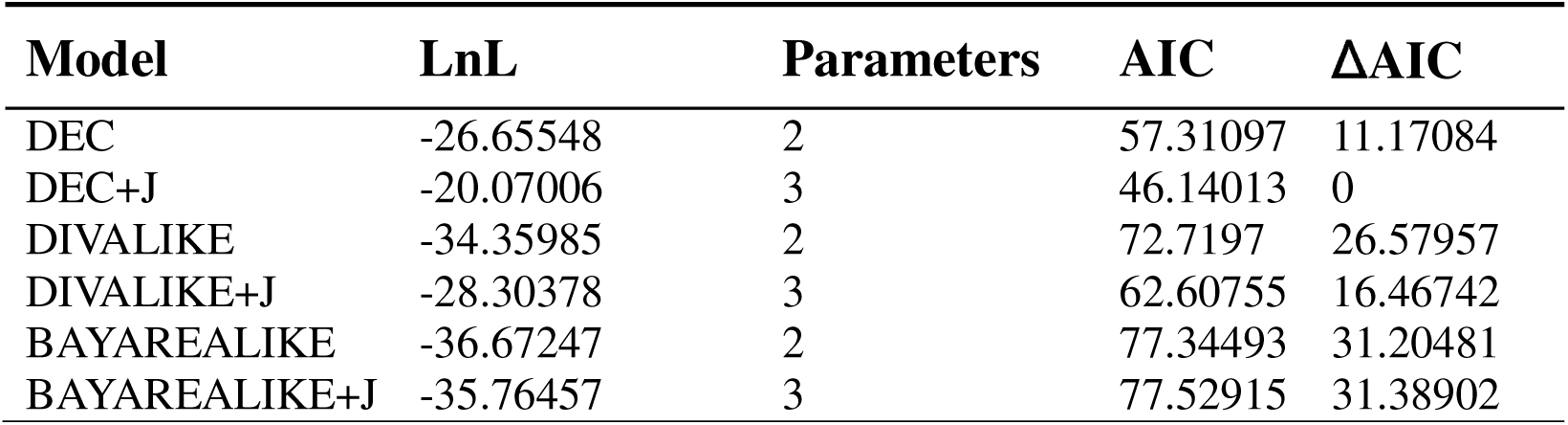
Statistics for the six models considered in the nuclear based analysis in BioGeoBEARS.

**Table S1.**
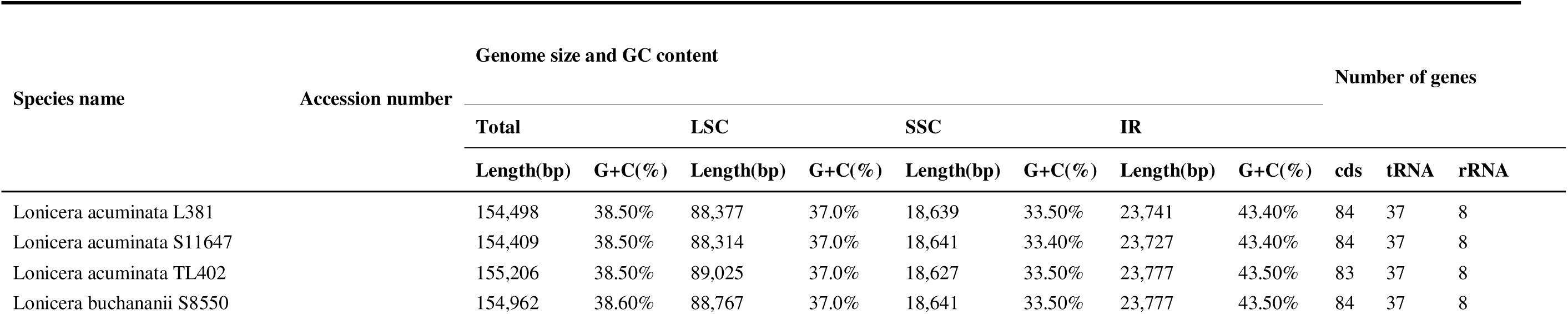
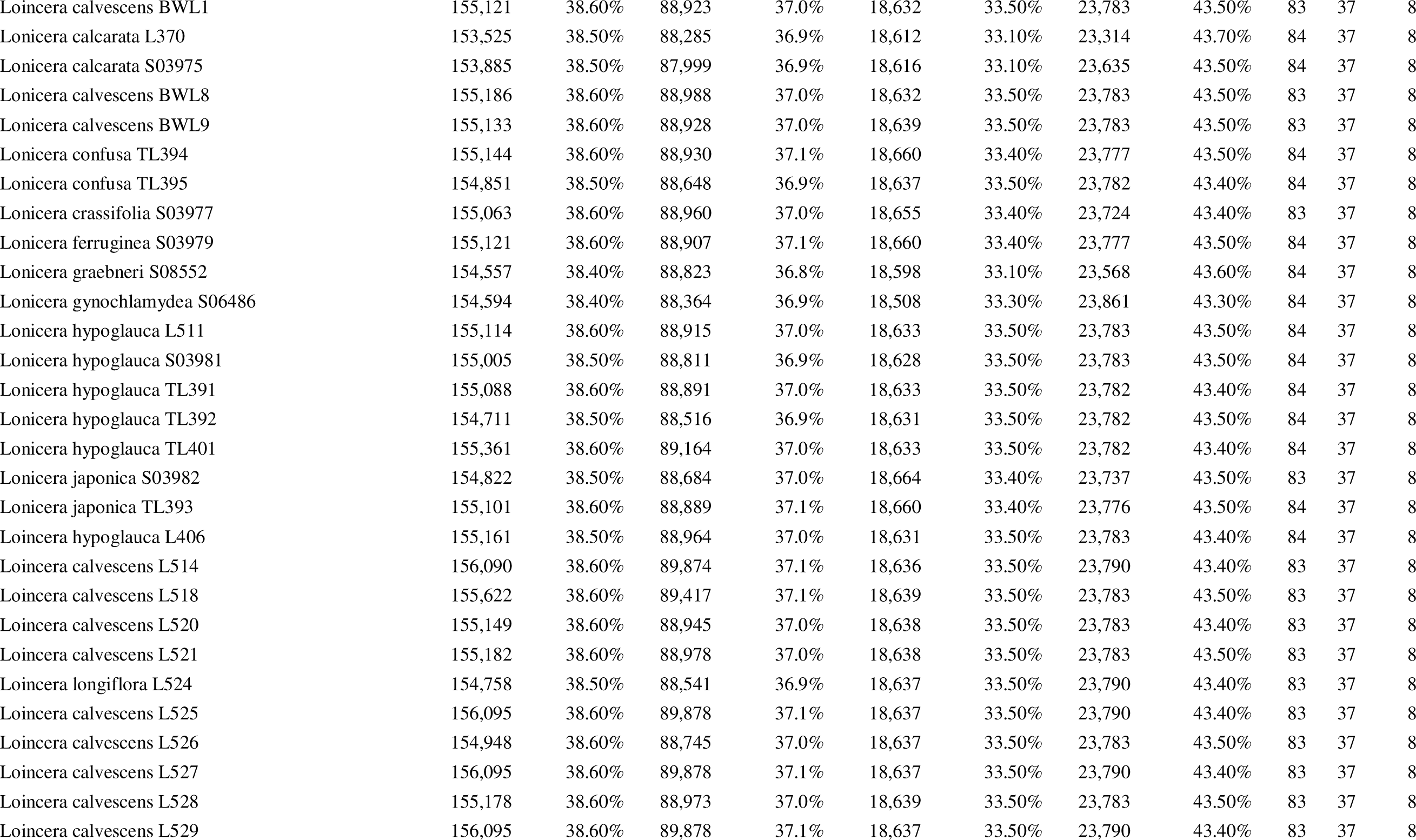
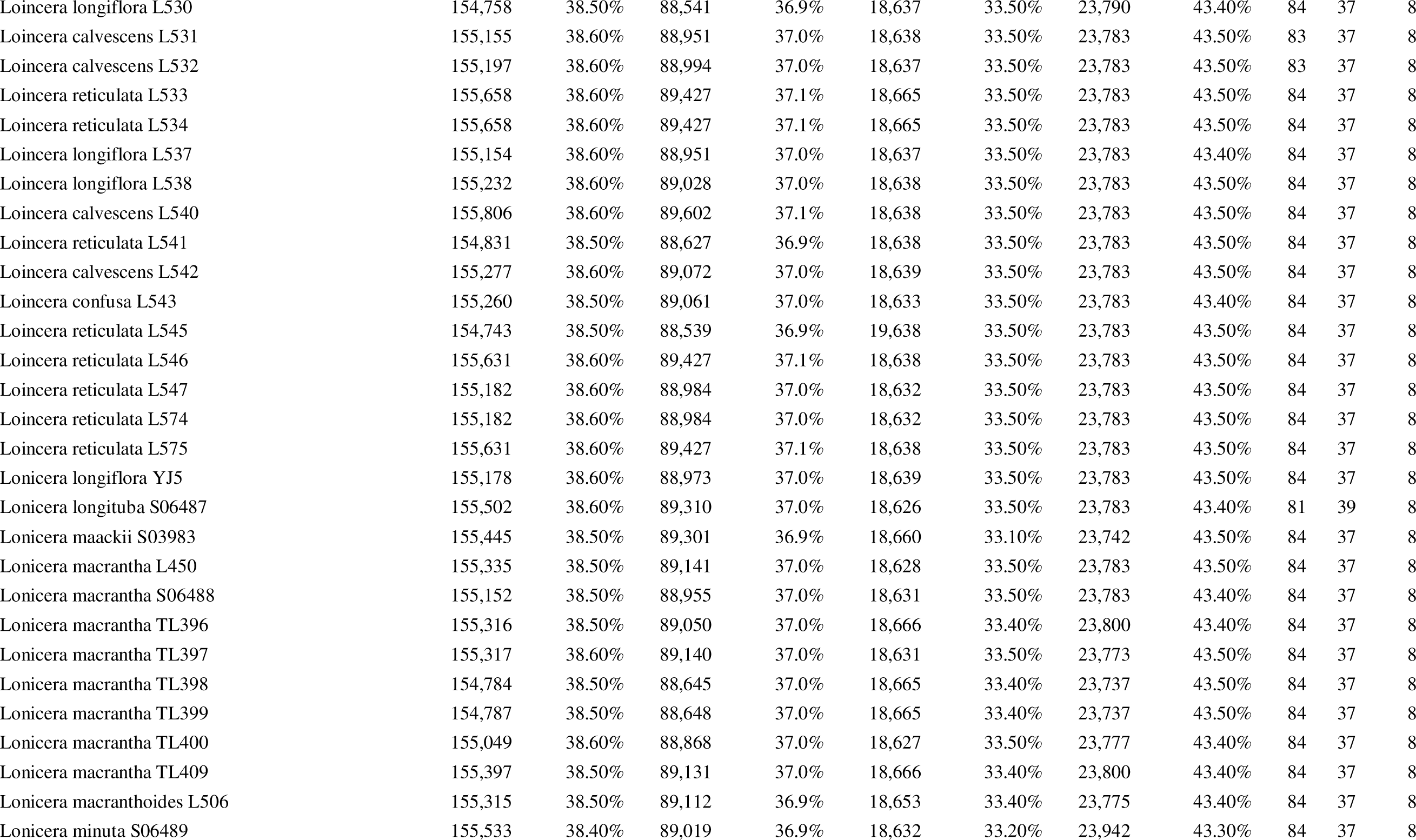
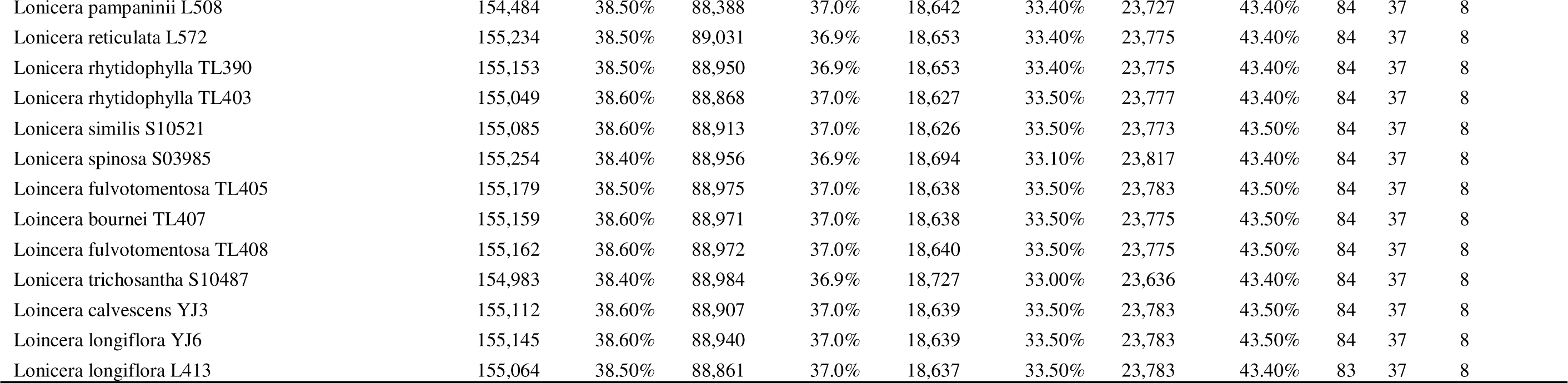
Length Information of Each Partition in Assembled Chloroplast Gene Data.

The results from the DEC+J model suggested that *Lonicera* from Hainan Island and Vietnam originated from mainland China (Figure 6). This model showed that the ancestors were likely distributed across these regions, with several long-distance dispersal events leading to the current distributions.

Similarly, the RASP analysis supported the findings from BioGeoBEARS(Figure S5). Each sample was assigned to its respective region based on its contemporary distribution range, corroborating the hypothesis that *Lonicera* species in Hainan Island and Vietnam have origins in mainland China and underwent four dispersal events and four vicariance events, shaping their current distribution patterns.

## 4. Discussion

### 4.1 Discordance among individual gene trees and species trees

Gene tree incongruence is common in phylogenetic studies, with multiple factors contributing to this phenomenon such as stochastic noise from uninformative genes, systematic errors, incomplete lineage sorting (ILS), hybridization/gene flow, and horizontal gene transfer. These factors can act independently or in combination, especially in lineages that have undergone rapid radiations (Knowles et al., 2018; Meyer et al., 2016; Morales-Briones et al., 2021; Pease et al., 2016; Schrempf and Szöll\Hosi, 2020). A recent simulation study by Molloy and Warnow (2018) demonstrated that the levels of ILS and gene tree estimation error (GTEE) differentially impact the accuracy of species tree estimation methods, including concatenation, site-based concatenation, and summary methods. Previous studies have shown that discrepancies can arise when constructing phylogenetic trees using different datasets or methods, often due to gene flow complicating genetic information exchange between species (Hodel et al., 2022; Sun et al., 2022). In our study, we observed discrepancies between phylogenetic trees constructed from nuclear and chloroplast data. These inconsistencies are likely due to gene introgression, hybridization, and rapid radiation rather than differing analytical methods.

In recent years, several analytical methods have been developed to account for the factors leading to incongruence (Edwards et al., 2016; Mirarab et al., 2014; Wen and Nakhleh, 2018; Xu and Yang, 2016; Zhang et al., 2024; Zhu et al., 2019). Employing multiple methods is crucial for understanding evolutionary processes and achieving accurate phylogenetic reconstruction. Previous molecular phylogenetic studies of *Lonicera* mainly relied on chloroplast gene data (Theis et al., 2008; Wang et al., 2020; Yang et al., 2024). However, chloroplast is maternally inherited in most angiosperms and does not fully reflect the true evolutionary history given its uniparental inheritance. Additionally, while chloroplast genome data is valuable for phylogenetic tree construction, challenges such as chloroplast capture still exist, which can lead to incongruences between gene trees and conflicts between nuclear and cytoplasmic genomes (Nakaji et al., 2015). Previous studies have provided evidence that hybridization plays a significant role in the formation of *Lonicera* (Landrein et al., 2012; Wang et al., 2020).

Sun et al. (2023) noted the presence of gene flow between the ancestors of subfamilies and other groups within *Lonicera*. This gene flow may be an important pattern for the survival and development of temperate plants(Cornille et al., 2015; Mitchell et al., 2009; O’Donnell et al., 2021; Stankowski et al., 2019). To further investigate whether the nuclear-cytoplasmic conflicts are due to gene flow, we used PhyloNet to analyze reticulate events, inferring networks that reflect gene flow, gene introgression, and other complex evolutionary events, thereby capturing the intricate hybridization phenomena more accurately (Hodel et al., 2022).

By comparing phylogenetic trees constructed from nuclear and chloroplast data, we observed inconsistencies in the topologies among species within *Lonicera*. For example, *L. confusa* L543 clustered with the Hainan population in both the concatenated tree and the species tree, while in the plastid tree, *L. confusa* L543 formed a branch with *L. longituba* S06487 from mainland China (BS=74). Similarly, the positions of *Lonicera* from Vietnam varied in the different phylogenetic trees. In the species tree, Vietnamese *Lonicera* species clustered together, whereas in the concatenated tree, *L. macrantha* L450 and *L. fulvotomentosa* L405 were nested among *Lonicera* from mainland China. These diverse topologies suggest a complex evolutionary history. Our dataset of 491 orthologous genes strongly supported the monophyly of *Nintooa*. However, we did not resolve the interspecific relationships within *Nintooa*, with different species often clustering together with low support for *Lonicera* from Hainan, indicating rapid diversification in this region. The network(Figure 5) evolution analysis also showed that *Nintooa* is a complex group with extensive hybridization and gene flow both within and between different sections. These gene flow events are one major cause of conflict between chloroplast and nuclear phylogenetic trees. The morphological similarity and overlapping habitats of species within *Lonicera*, such as the *Lonicera macrantha* complex, provide conditions for hybridization, suggesting a complex origin and evolutionary process for *Lonicera*.

### 4.2 Systematic position of subsection *Calcaratae*

In this study, we resolved the phylogenetic relationships of Section *Nintooa*. Both the coalescent and concatenated tree strongly support Section *Nintooa* as the sister group to Subgenus *Chamaecerasus* (Figures 2 and 3) Our results align with those of Srivastav et al. (2023).. In contrast, previous analyses placed most Section *Nintooa* species within a well-supported nested clade of Subgenus *Chamaecerasus*, with a sister clade composed of *Isoxylosteum*, *Coeloxylosteum*, and *Isika*.

Rehder (1903) classified *L. calcarata* within Calcaratae. Previous analyses separated *L. calcarata* from other Section Nintooa species (Nakaji et al., 2015; Theis et al., 2008). Our findings support this view; however, different methods show varying phylogenetic positions for *L. calcarata*. In both the species and concatenated trees, *L. calcarata* formed a sister group with all other Section Nintooa species. Conversely, in the chloroplast genome data, *L. calcarata* was sister to Subgenus *Chamaecerasus*. Both chloroplast and nuclear gene data robustly support a distinct phylogenetic position for Calcaratae, indicating a potential true nuclear-cytoplasmic conflict(LeeLYaw et al., 2019).

Unlike other Nintooa species, *L. calcarata* exhibits fused bracts, fused ovaries, and long nectar spurs(Srivastav et al., 2023). Network analyses also indicate that the formation of *Calcaratae* involved hybridization events with *Nintooa* and gene flow with ancestors of other species within Subgenus *Chamaecerasus*. Additionally, Hu et al. (2008) found that the plastid inheritance in *Lonicera* might be biparental, suggesting a complex cause for the nuclear-cytoplasmic conflict in *Calcaratae*.

The hypothesis of *L. calcarata* being phylogenetically distinct within *Nintooa* due to its unique morphological traits and hybridization events is more plausible. The fusion of bracts and ovaries, along with long nectar spurs, supports the hypothesis that *L. calcarata* has undergone significant divergence. This, combined with evidence of gene flow and potential biparental plastid inheritance, indicates that *L. calcarata* occupies a unique evolutionary niche within *Nintooa*.

### 4.3 Divergence Time and Ancestral Distribution Reconstruction

Using nuclear data and two fossil calibration points, we estimated the divergence times of subgenus *Lonicera*. Our results indicate that the common ancestor of *Lonicera* dates to the mid-Eocene, 42.55 Ma (95% HPD: 31.44–52.11 Ma). The oldest *Lonicera* species are from the continental region, with the divergence times of the three major continental clades ranging from 17.41 to 20.04 Ma. The most recent common ancestor of the Hainan and Vietnam *Lonicera* species is estimated to be 15.74 Ma (95% HPD: 11.58–19.28 Ma), with the divergence of Hainan *Lonicera* dating to 14.04 Ma (95% HPD: 10.62–17.04 Ma) and the Vietnam species to 14.15 Ma (95% HPD: 9.89–17.69 Ma). Sun and Li (2003) hypothesized that the disappearance of the Tethys Sea and the uplift of the Himalayas, were driving factors preserving *Lonicera* species, which subsequently dispersed across the Hengduan Mountains and the Qinghai-Tibet Plateau.

In this study, *Lonicera* experienced a rapid radiation from the mid-Oligocene to the late Miocene, with major lineages diversifying within a short period. The diversification of continental *Lonicera* species mainly occurred between 26.45 and 17.09 Ma, coinciding with several significant geological events, including the second uplift of the Qinghai-Tibet Plateau around 21 Ma. This period was characterized by the westward retreat of the Paratethys Sea and increased aridity in inland Asia. The diversification of Hainan *Lonicera* species occurred during the Miocene (5.3–23 Ma), corresponding to significant geological events on Hainan Island. The Wuzhishan region of Hainan experienced two rapid uplift and denudation phases. The first phase occurred from the Oligocene to the Miocene (32–17 Ma), characterized by the subduction and retreat of the Pacific Plate beneath the Eurasian Plate and the second expansion of the South China Sea, leading to rapid uplift in the Qiongzhong Mountains until the mid-Miocene. The second phase of rapid uplift and erosion occurred at the end of the Miocene (5 Ma), with the end of the South China Sea expansion and the subduction of the Philippine Plate beneath the Asian Plate, leading to accelerated thermal subsidence and drastic climate change, causing extensive uplift and accelerated erosion of Hainan Island (Shi et al., 2011; Shi et al.,2024). These geological and climatic changes profoundly impacted vegetation and ecosystems. *Lonicera* likely evolved rapidly in response to these environmental changes, adapting to new ecological conditions. This highlights the interaction of plants with their environment throughout geological history, resulting in the evolution of diverse species to adapt to different ecological niches.

Our ancestral distribution reconstruction results indicate that both Hainan Island and continental *Lonicera* originated from mainland China. Our findings support the hypothesis of an East Asian origin. However, due to the lack of more comprehensive samples from other continental regions, further research is needed.

## 5 Conclusions

The analyses of chloroplast and nuclear data yielded inconsistent results with *Section Nintooa* forming a monophyletic group that is sister to other species within the Caprifoliaceae. *Volubilis* was monophyletic and sister to *Calcaratae* in both the species tree and concatenated nuclear trees. However, in the chloroplast tree, *Calcaratae* is sister to all other *Subgenus Chamaecerasus* species. Additionally, *Lonicera* from Hainan Island show low support and short branch lengths in the chloroplast tree, indicating that chloroplast data does not provide enough signal suggesting a rapid radiation. Although molecular evidence does not clearly differentiate species of Section *Nintooa*, the lack of differentiation supports the *Lonicera macrantha* complex hypothesis.

We found that *Lonicera* from Hainan Island are sister to those from Vietnam, suggesting historical connections between Hainan and Vietnam. Divergence time estimates indicate that most Section *Nintooa* species originated during the Pliocene to Miocene, specifically between 5-14 Ma. Our biogeographic analysis supports the ancestor of Section *Nintooa* originated in mainland China, underwent long-distance dispersal and island drift, and experienced a series of complex geological events that shaped the current distribution of Section *Nintooa*. This study better resolves the deep phylogenetic relationships among the main lineages within Section *Nintooa*.

## Author contributions

H.F.W. conceived the study. HFW, SYZ, HYL and HJD performed the research, collect samples and analyzed the data. S.Y.Z., Q.H.S., D.F.M-B., H.X.W., J.W., J.B.L, and H.F.W. wrote and revised the manuscript.

## Acknowledgements

The authors appreciate the constructive comments and suggestions from the editor and reviewers on our initial manuscript. The work was funded by National Science Foundation of China (32270221), the Project of Sanya Yazhou Bay Science and Technology City (Grant number: SCKJ-JYRC-2022-83), and Hainan Provincial Natural Science Foundation of China (421RC486 and 822QN314). We thank Gabriel Johnson for his assistance with the target enrichment experiment, and the United States National Herbarium for collection access. We acknowledge the staff in the Laboratories of Analytical Biology at the National Museum of Natural History, the Smithsonian Institution for technical support and assistance.

---

**Figure S1.** Phylogeny of *Lonicera* inferred using Astral. Support values displayed on the branches are Quartet Concordance (QC)/Quartet Differential (QD)/Quartet Informativeness (QI).

**Figure S2.** A maximum likelihood cladogram of Subgenus *Chamaecerasus* inferred with IQ-tree using the nuclear concatenated matrix. Support values displayed on the branches are Quartet Concordance (QC)/Quartet Differential (QD)/Quartet Informativeness (QI)

**Figure S3.** A maximum likelihood cladogram of Subgenus *Chamaecerasus* inferred with IQ-tree using the plastome matrix. Support values displayed on the branches are Quartet Concordance (QC)/Quartet Differential (QD)/Quartet Informativeness (QI).

**Figure S4.** The DIVALIKE model in BioGeoBEARS was utilized to infer the ancestral range of Section Nintooa. The geographic distribution areas are delineated as follows: (A) Hainan Island; (B) Mainland China; (C) Vietnam.

**Figure S5.** The S-DIVALIKE model in RASP was utilized to infer the ancestral range of Section Nintooa. The geographic distribution areas are delineated as follows: (A) Hainan Island; (B) Mainland China; (C) Vietnam.

**Figure S6.** Phylogenetic tree constructed based on the large single-copy region of the plastome.

**Figure S7.** Phylogenetic tree constructed based on the small single-copy region of the plastome.

**Figure S8.** Pphylogenetic tree based on the removal of the inverted repeat region of the plastome.

**Figure S9.** Phylogenetic tree constructed based on the non-coding region of the complete plastome.

**Figure S10.** Phylogenetic tree constructed based on coding regions of the complete plastome.

**Figure S11.** Phylogenetic tree constructed based on the one Inverted Repeat.

## References

Baker, W.J., Dodsworth, S., Forest, F., Graham, S.W., Johnson, M.G. (TTU), McDonnell, A., Pokorny, L., Tate, J.A., Wicke, S., Wickett, N.J., 2021. Exploring Angiosperms353: An open, community toolkit for collaborative phylogenomic research on flowering plants.

Bellard, C., Leclerc, C., Courchamp, F., 2014. Impact of sea level rise on the 10 insular biodiversity hotspots. Global Ecology and Biogeography 23, 203–212.

Bouckaert, R., Heled, J., Kühnert, D., Vaughan, T., Wu, C.-H., Xie, D., Suchard, M.A., Rambaut, A., Drummond, A.J., 2014. BEAST 2: A Software Platform for Bayesian Evolutionary Analysis. PLoS Comput Biol 10, e1003537.

Brown, J.W., Walker, J.F., Smith, S.A., 2017. Phyx: phylogenetic tools for unix. Bioinformatics 33, 1886–1888.

Chen, D., Chang, J., Li, S.-H., Liu, Y., Liang, W., Zhou, F., Yao, C.-T., Zhang, Z., 2015. Was the exposed continental shelf a long-distance colonization route in the ice age? The Southeast Asia origin of Hainan and Taiwan partridges. Molecular Phylogenetics and Evolution 83, 167–173.

Chen, D., Zhang, X., Kang, H., Sun, X., Yin, S., Du, H., Yamanaka, N., Gapare, W., Wu, H.X., Liu, C., 2012. Phylogeography of Quercus variabilis based on chloroplast DNA sequence in East Asia: multiple glacial refugia and mainland-migrated island populations. PLoS One 7, e47268.

Cornille, A., Feurtey, A., Gélin, U., Ropars, J., Misvanderbrugge, K., Gladieux, P., Giraud, T., 2015. Anthropogenic and natural drivers of gene flow in a temperate wild fruit tree: a basis for conservation and breeding programs in apples. Evolutionary applications 8, 373–384.

Crowl, A.A., Visger, C.J., Mansion, G., Hand, R., Wu, H., Kamari, G., Phitos, D., Cellinese, N., 2015. Evolution and biogeography of the endemic Roucela complex (Campanulaceae: Campanula) in the Eastern Mediterranean. Ecology and Evolution 5, 5329–5343.

Darling, A.C.E., Mau, B., Blattner, F.R., Perna, N.T., 2004. Mauve: Multiple Alignment of Conserved Genomic Sequence With Rearrangements. Genome Res. 14, 1394– 1403.

Doyle, J.J., Doyle, J.L., 1987. A rapid DNA isolation procedure for small quantities of fresh leaf tissue. Phytochemical Bulletin 19, 11–15.

Drummond, A.J., Suchard, M.A., Xie, D., Rambaut, A., 2012. Bayesian Phylogenetics with BEAUti and the BEAST 1.7. Molecular Biology and Evolution 29, 1969–1973.

Edwards, S.V., Xi, Z., Janke, A., Faircloth, B.C., McCormack, J.E., Glenn, T.C., Zhong, B., Wu, S., Lemmon, E.M., Lemmon, A.R., Leaché, A.D., Liu, L., Davis, C.C., 2016. Implementing and testing the multispecies coalescent model: A valuable paradigm for phylogenomics. Molecular Phylogenetics and Evolution 94, 447–462.

Friis, E.M., 1985. Angiosperm fruits and seeds from the Middle Miocene of Jutland (Denmark), Biologiske Skrifter. The Royal Danish Academy of Sciences and Letters, Copenhagen.

Hara, H., 1983. A revision of Caprifoliaceae of Japan with reference to allied plants in other districts and the Adoxaceae. Ginkgoana No.5. Academia Scientific Book Inc., Tokyo.

Hodel, R.G.J., Zimmer, E.A., Liu, B.-B., Wen, J., 2022. Synthesis of Nuclear and Chloroplast Data Combined With Network Analyses Supports the Polyploid Origin of the Apple Tribe and the Hybrid Origin of the Maleae—Gillenieae Clade. Front. Plant Sci. 12.

Hong-Jin, D., Hua, P., 2014. Relationships within the Lonicera macrantha Complex Based on Morphological and Molecular Data. Plant Diversity 36, 133–141.

Hsu, P.S., Wang, H.J., 1988. Lonicera, in: Flora Reipublicae Popularis Sinicae. Science Press, Beijing, pp. 143–259, pls. 38–69.

Hu, Y., Zhang, Q., Rao, G., Sodmergen, 2008. Occurrence of Plastids in the Sperm Cells of Caprifoliaceae: Biparental Plastid Inheritance in Angiosperms is Unilaterally Derived from Maternal Inheritance. Plant and Cell Physiology 49, 958–968.

Huang, P.-H., Wang, T.-R., Li, M., Lu, Z.-J., Su, R.-P., Fang, O.-Y., Li, L., Zhou, S.-S., Tan, Y.-H., Meng, H.-H., Song, Y.-G., Li, J., 2024. RAD-seq data for Engelhardia roxburghiana provide insights into the palaeogeography of Hainan Island and its relationship to mainland China since the late Eocene. Palaeogeography, Palaeoclimatology, Palaeoecology 651, 112392.

Huson, D.H., Scornavacca, C., 2012. Dendroscope 3: An Interactive Tool for Rooted Phylogenetic Trees and Networks. Systematic Biology 61, 1061–1067.

Jin, J.-J., Yu, W.-B., Yang, J.-B., Song, Y., dePamphilis, C.W., Yi, T.-S., Li, D.-Z., 2020. GetOrganelle: a fast and versatile toolkit for accurate de novo assembly of organelle genomes. Genome Biology 21, 241.

Katoh, K., Standley, D.M., 2013. MAFFT Multiple Sequence Alignment Software Version 7: Improvements in Performance and Usability. Molecular Biology and Evolution 30, 772–780.

Kearse, M., Moir, R., Wilson, A., Stones-Havas, S., Cheung, M., Sturrock, S., Buxton, S., Cooper, A., Markowitz, S., Duran, C., Thierer, T., Ashton, B., Meintjes, P., Drummond, A., 2012. Geneious Basic: an integrated and extendable desktop software platform for the organization and analysis of sequence data. Bioinformatics 28, 1647–1649.

Knowles, L.L., Huang, H., Sukumaran, J., Smith, S.A., 2018. A matter of phylogenetic scale: Distinguishing incomplete lineage sorting from lateral gene transfer as the cause of gene tree discord in recent versus deep diversification histories. American J of Botany 105, 376–384.

Lańcucka-Rodoniowa, M., 1967. Two new genera: Hemiptelea Planch. and Weigela Thumb. in the younger tertiary of Poland. Acta Palaeobotanica 8, 1–17.

Landis, M., Matzke, N., Huelsenbeck, J., Moore, B., 2013. Bayesian analysis of biogeography when the number of areas is large. Systematic Biology 62.

Landrein, S., Prenner, G., Chase, M.W., Clarkson, J.J., 2012. Abelia and relatives: phylogenetics of Linnaeeae (Dipsacales–Caprifoliaceae sl) and a new interpretation of their inflorescence morphology. Botanical journal of the Linnean Society 169, 692–713.

Lee, J.-H., Lee, D.-H., Choi, B.-H., 2013. Phylogeography and genetic diversity of East Asian Neolitsea sericea (Lauraceae) based on variations in chloroplast DNA sequences. J Plant Res 126, 193–202.

Lee-Yaw, J.A., Grassa, C.J., Joly, S., Andrew, R.L., Rieseberg, L.H., 2019. An evaluation of alternative explanations for widespread cytonuclear discordance in annual sunflowers ( Helianthus ). New Phytologist 221, 515–526.

Li, H.T., Yi, T.S., Gao, L.M., Ma, P.F., Zhang, T., Yang, J.B., Gitzendanner, M.A., Fritsch, P.W., Cai, J., Luo, Y., Wang, H., Bank, M.V., Zhang, S.D., Wang, Q.F., Wang, J., Zhang, Z.R., Fu, C.N., Yang, J., Hollingsworth, P.M., Chase, M.W., Soltis, D.E., Soltis, P.S., Li, D.Z., 2019. Origin of angiosperms and the puzzle of the Jurassic gap. Nature Plants 5, 461–470.

Lin, A.-Q., Csorba, G., Li, L.-F., Jiang, T.-L., Lu, G.-J., Thong, V.D., Soisook, P., Sun, K.- P., Feng, J., 2014. Phylogeography of Hipposideros armiger (Chiroptera: Hipposideridae) in the Oriental Region: the contribution of multiple Pleistocene glacial refugia and intrinsic factors to contemporary population genetic structure. Journal of Biogeography 41, 317–327.

Mai, U., Mirarab, S., 2018. TreeShrink: fast and accurate detection of outlier long branches in collections of phylogenetic trees. BMC Genomics 19, 272.

Matzke, N.J., 2014. Model Selection in Historical Biogeography Reveals that Founder-Event Speciation Is a Crucial Process in Island Clades. Systematic Biology 63, 951–970.

Matzke, N.J., 2013. Probabilistic historical biogeography: new models for founder-event speciation, imperfect detection, and fossils allow improved accuracy and model-testing. Frontiers of Biogeography 5.

Maximowicz, C.J., 1877. Diagnoses plantarum novarum Asiaticarum. Imperialis Academiae Scientiarum, Petropli.

Meyer, B.S., Matschiner, M., Salzburger, W., 2016. Disentangling Incomplete Lineage Sorting and Introgression to Refine Species-Tree Estimates for Lake Tanganyika Cichlid Fishes. Syst Biol syw069.

Mirarab, S., Bayzid, Md.S., Boussau, B., Warnow, T., 2014. Statistical binning enables an accurate coalescent-based estimation of the avian tree. Science 346, 1250463.

Mitchell, R.J., Irwin, R.E., Flanagan, R.J., Karron, J.D., 2009. Ecology and evolution of plant–pollinator interactions. Annals of Botany 103, 1355–1363.

Molloy, E.K., Warnow, T., 2018. To Include or Not to Include: The Impact of Gene Filtering on Species Tree Estimation Methods. Systematic Biology 67, 285–303.

Morales-Briones, D.F., Kadereit, G., Tefarikis, D.T., Moore, M.J., Smith, S.A., Brockington, S.F., Timoneda, A., Yim, W.C., Cushman, J.C., Yang, Y., 2021. Disentangling Sources of Gene Tree Discordance in Phylogenomic Data Sets: Testing Ancient Hybridizations in Amaranthaceae s.l. Systematic Biology 70, 219–235.

Myers, N., Mittermeier, R.A., Mittermeier, C.G., Da Fonseca, G.A., Kent, J., 2000. Biodiversity hotspots for conservation priorities. Nature 403, 853–858.

Nakai, T., 1938. A new classification of the genus Lonicera in the Japanese empire, together with the diagnoses of new species and new varieties. J. Jap. Bot. 14, 359–376.

Nakaji, M., Tanaka, N., Sugawara, T., 2015. A Molecular Phylogenetic Study of Lonicera L. (Caprifoliaceae) in Japan Based on Chloroplast DNA Sequences 66.

O’Donnell, S.T., Fitz-Gibbon, S.T., Sork, V.L., 2021. Ancient introgression between distantly related white oaks (Quercus sect. Quercus) shows evidence of climate-associated asymmetric gene exchange. Journal of Heredity 112, 663–670.

Ortiz, E.M., Hoewener, A., Shigita, G., Raza, M., Maurin, O., Zuntini, A., Forest, F., Baker, W.J., Schaefer, H., 2023. A novel phylogenomics pipeline reveals complex pattern of reticulate evolution in Cucurbitales. bioRxiv 2023–10.

Pease, J.B., Brown, J.W., Walker, J.F., Hinchliff, C.E., Smith, S.A., 2018. Quartet Sampling distinguishes lack of support from conflicting support in the green plant tree of life. American J of Botany 105, 385–403.

Pease, J.B., Haak, D.C., Hahn, M.W., Moyle, L.C., 2016. Phylogenomics Reveals Three Sources of Adaptive Variation during a Rapid Radiation. PLoS Biol 14, e1002379.

Qiu, Y.-X., Fu, C.-X., Comes, H.P., 2011. Plant molecular phylogeography in China and adjacent regions: tracing the genetic imprints of Quaternary climate and environmental change in the world’s most diverse temperate flora. Molecular phylogenetics and evolution 59, 225–244.

Rambaut, A., Drummond, A.J., Xie, D., Baele, G., Suchard, M.A., 2018. Posterior summarisation in Bayesian phylogenetics using Tracer 1.7. Systematic biology 67.

Ree, R.H., Smith, S.A., 2008. Maximum Likelihood Inference of Geographic Range Evolution by Dispersal, Local Extinction, and Cladogenesis. Systematic Biology 57, 4–14.

Rehder, A., 1903. Synopsis of the genus Lonicera. Ann. Rep. Missouri Bot. Gard. 14, 27– 232.

Ronquist, F., 1997. Dispersal-Vicariance Analysis: A New Approach to the Quantification of Historical Biogeography. Systematic Biology 46, 195.

Sayyari, E., Mirarab, S., 2016. Fast Coalescent-Based Computation of Local Branch Support from Quartet Frequencies. Mol Biol Evol 33, 1654–1668.

Schrempf, D., Szöll-Hosi, G., 2020. The sources of phylogenetic conflicts. Phylogenetics in the genomic era 3–1.

Shi, J., Han, S., Du, J., Han, J., Sun, D., Hu, D., 2024. Surface process response to the expansion of the northern South China Sea: Age evidence of apatite fission tracks from Wuzhi Mountains, Hainan Island. Acta Geologica Sinica 98, 421–432.

Shi, X., Kohn, B., Spencer, S., Guo, X., Li, Y., Yang, X., Shi, H., Gleadow, A., 2011. Cenozoic denudation history of southern Hainan Island, South China Sea: Constraints from low temperature thermochronology. Tectonophysics 504, 100– 115.

Srivastav, M., Clement, W.L., Landrein, S., Zhang, J., Howarth, D.G., Donoghue, M.J., 2023. A phylogenomic analysis of Lonicera and its bearing on the evolution of organ fusion. American J of Botany 110, e16143.

Stamatakis, A., 2014. RAxML version 8: a tool for phylogenetic analysis and post-analysis of large phylogenies. Bioinformatics 30, 1312–1313.

Stankowski, S., Chase, M.A., Fuiten, A.M., Rodrigues, M.F., Ralph, P.L., Streisfeld, M.A., 2019. Widespread selection and gene flow shape the genomic landscape during a radiation of monkeyflowers. PLoS biology 17, e3000391.

Sun, Q.-H., Morales-Briones, D.F., Wang, H.-X., Landis, J.B., Wen, J., Wang, H.-F., 2023. Target sequence capture data shed light on the deeper evolutionary relationships of subgenus Chamaecerasus in Lonicera (Caprifoliaceae). Molecular Phylogenetics and Evolution 184, 107808.

Sun, Q.-H., Morales-Briones, D.F., Wang, H.-X., Landis, J.B., Wen, J., Wang, H.-F., 2022. Phylogenomic analyses of the East Asian endemic Abelia (Caprifoliaceae) shed insights into the temporal and spatial diversification history with widespread hybridization. Annals of Botany 129, 201–216.

Tang, Z., Wang, Z., Zheng, C., Fang, J., 2006. Biodiversity in China’s mountains. Frontiers in Ecology and the Environment 4, 347–352.

Taylor, S., Kumar, L., 2016. Global Climate Change Impacts on Pacific Islands Terrestrial Biodiversity: A Review. Tropical Conservation Science 9, 203–223.

Theis, N., Donoghue, M.J., Li, J., 2008. Phylogenetics of the Caprifolieae and Lonicera (Dipsacales) Based on Nuclear and Chloroplast DNA Sequences. Systematic Botany 33, 776–783.

Tian, X., Wang, Q., Zhou, Y., 2018. Euphorbia Section Hainanensis (Euphorbiaceae), a New Section Endemic to the Hainan Island of China From Biogeographical, Karyological, and Phenotypical Evidence. Front. Plant Sci. 9, 660.

Voris, H.K., 2000. Maps of Pleistocene sea levels in Southeast Asia: shorelines, river systems and time durations. Journal of Biogeography 27, 1153–1167.

Wallace, A.R., 1869. Sir Charles Lyell on geological climates and the origin of species. Quarterly review 126, 359–94.

Wang, H.-X., Liu, H., Moore, M.J., Landrein, S., Liu, B., Zhu, Z.-X., Wang, H.-F., 2020. Plastid phylogenomic insights into the evolution of the Caprifoliaceae s.l. (Dipsacales). Molecular Phylogenetics and Evolution 142, 106641.

Wen, D., Nakhleh, L., 2018. Coestimating reticulate phylogenies and gene trees from multilocus sequence data. Systematic biology 67, 439–457.

Wick, R.R., Schultz, M.B., Zobel, J., Holt, K.E., 2015. Bandage: interactive visualization of de novo genome assemblies. Bioinformatics 31, 3350–3352.

Xu, B., Yang, Z., 2016. Challenges in Species Tree Estimation Under the Multispecies Coalescent Model. Genetics 204, 1353–1368.

Yang, Q., Landrein, S., 2011. Linnaeaceae, in: Wu, Z., Hong, D., Raven, P.H. (Eds.), Flora of China. Science Press and Missouri Botanical Garden Press, Beijing and St. Louis, pp. 642–648.

Yang, Q., Landrein, S., Osborne, J., Borosova, R., 2011. Caprifoliaceae. Flora of China 19, 616–641.

Yang, X., Sun, Q., Morales-Briones, D.F., Landis, J.B., Chen, D., Wang, Hong-Xin, Wen, J., Wang, Hua-Feng, 2024. New insights into infrageneric relationships of Lonicera (Caprifoliaceae) as revealed by nuclear ribosomal DNA cistron data and plastid phylogenomics. J of Sytematics Evolution 62, 333–357.

Yang, Y., Smith, S.A., 2014. Orthology Inference in Nonmodel Organisms Using Transcriptomes and Low-Coverage Genomes: Improving Accuracy and Matrix Occupancy for Phylogenomics. Molecular Biology and Evolution 31, 3081–3092.

Ye, J.-W., Zhang, Y., Wang, X.-J., 2017. Phylogeographic breaks and the mechanisms of their formation in the Sino-Japanese floristic region. Chinese Journal of Plant Ecology 41, 1003–1019.

Yu, Y., Harris, A.J., Blair, C., He, X., 2015. RASP (Reconstruct Ancestral State in Phylogenies): a tool for historical biogeography. Mol Phylogenet Evol 87, 46–49.

Yu, Y., Nakhleh, L., 2015. A maximum pseudo-likelihood approach for phylogenetic networks. BMC Genomics 16, S10.

Zhang, C., Rabiee, M., Sayyari, E., Mirarab, S., 2018. ASTRAL-III: polynomial time species tree reconstruction from partially resolved gene trees. BMC Bioinformatics 19, 153.

Zhang, D., Ye, Z., Yamada, K., Zhen, Y., Zheng, C., Bu, W., 2016. Pleistocene sea level fluctuation and host plant habitat requirement influenced the historical phylogeography of the invasive species Amphiareus obscuriceps (Hemiptera: Anthocoridae) in its native range. BMC Evol Biol 16, 174.

Zhang, W., Ding, Y., Cao, Y., Li, P., Yang, Y., Pang, X., Bai, W., Zhang, D., 2024. Uncovering ghost introgression through genomic analysis of a distinct eastern Asian hickory species. The Plant Journal tpj.16859.

Zhu, H., 2017. Floristic characteristics and affinities in Lao PDR, with a reference to the biogeography of the Indochina peninsula. PLoS One 12, e0179966.

Zhu, H., 2016. Biogeographical Evidences Help Revealing the Origin of Hainan Island. PLoS ONE 11, e0151941.

Zhu, J., Liu, X., Ogilvie, H.A., Nakhleh, L.K., 2019. A divide-and-conquer method for scalable phylogenetic network inference from multilocus data. Bioinformatics 35, i370–i378.

