## Supplementary figures and images for "Target enrichment phylogenomics and biogeographic analyses unravel rapid radiation and reticulate evolution between Hainan-South China mainland -Vietnam in Section *Nintooa* (*Lonicera*, Caprifoliaceae)"

### Supplemental Figure S1

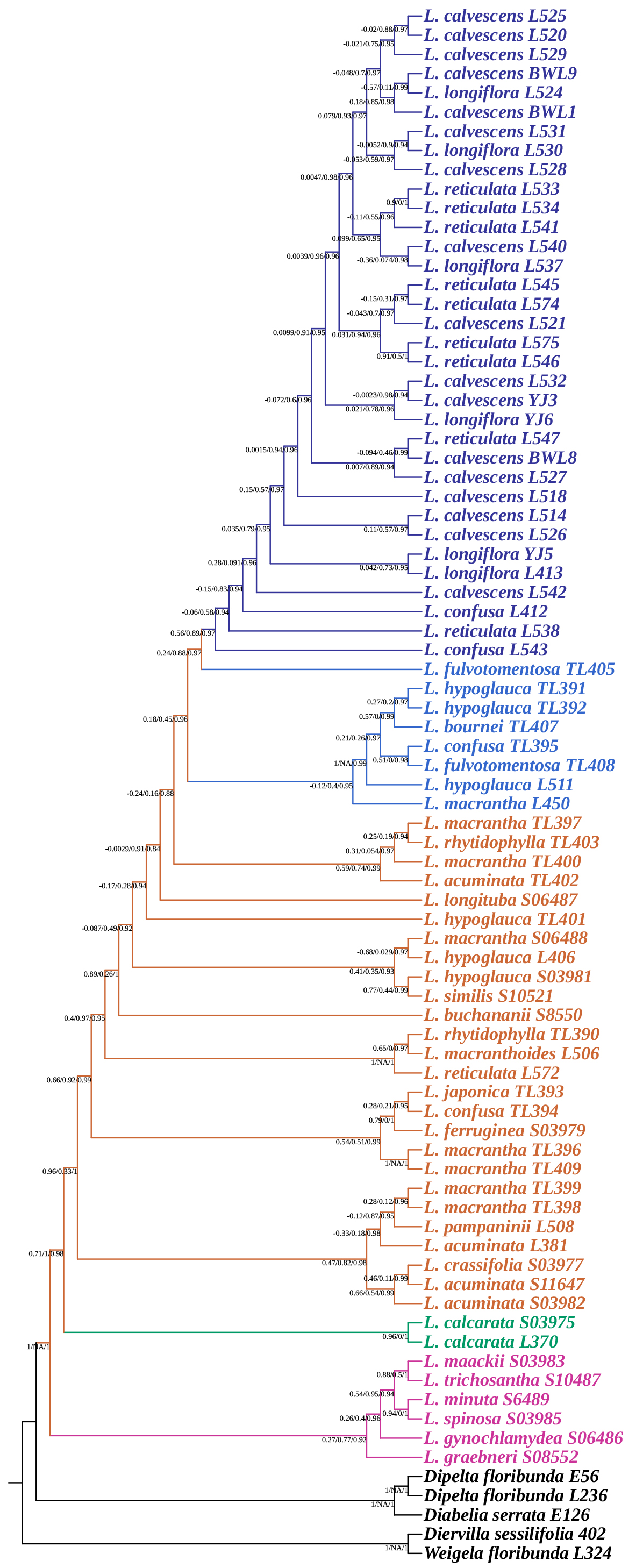

### Supplemental Figure S2

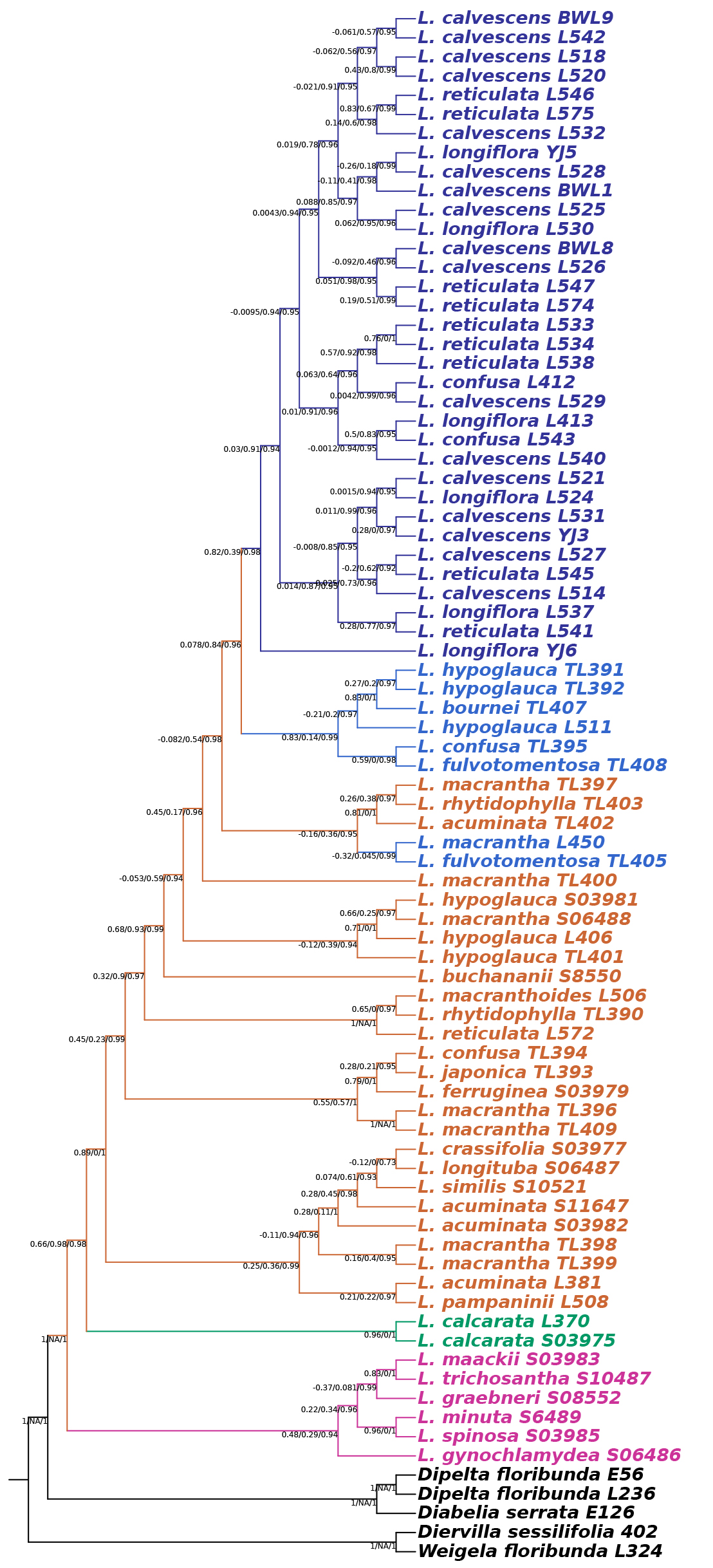

### Supplemental Figure S3

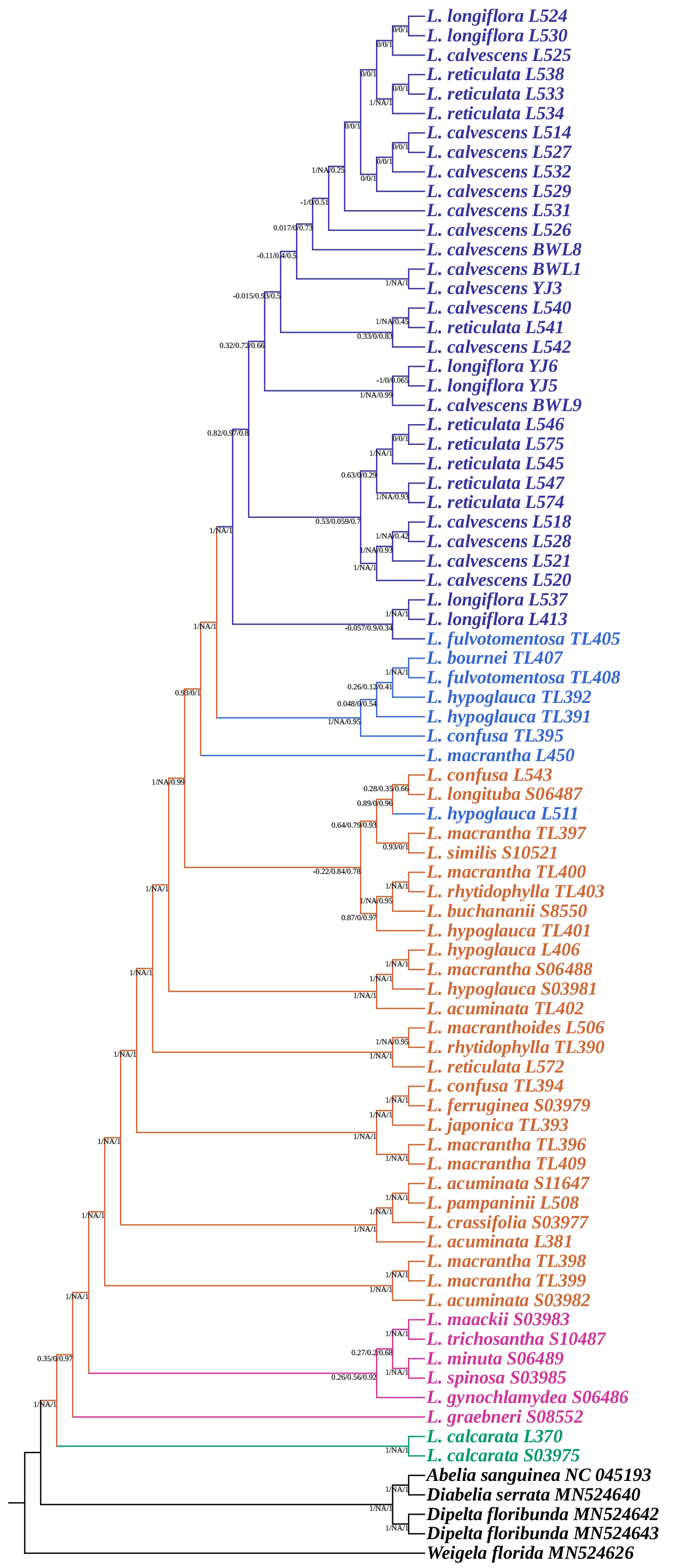

### Supplemental Figure S4

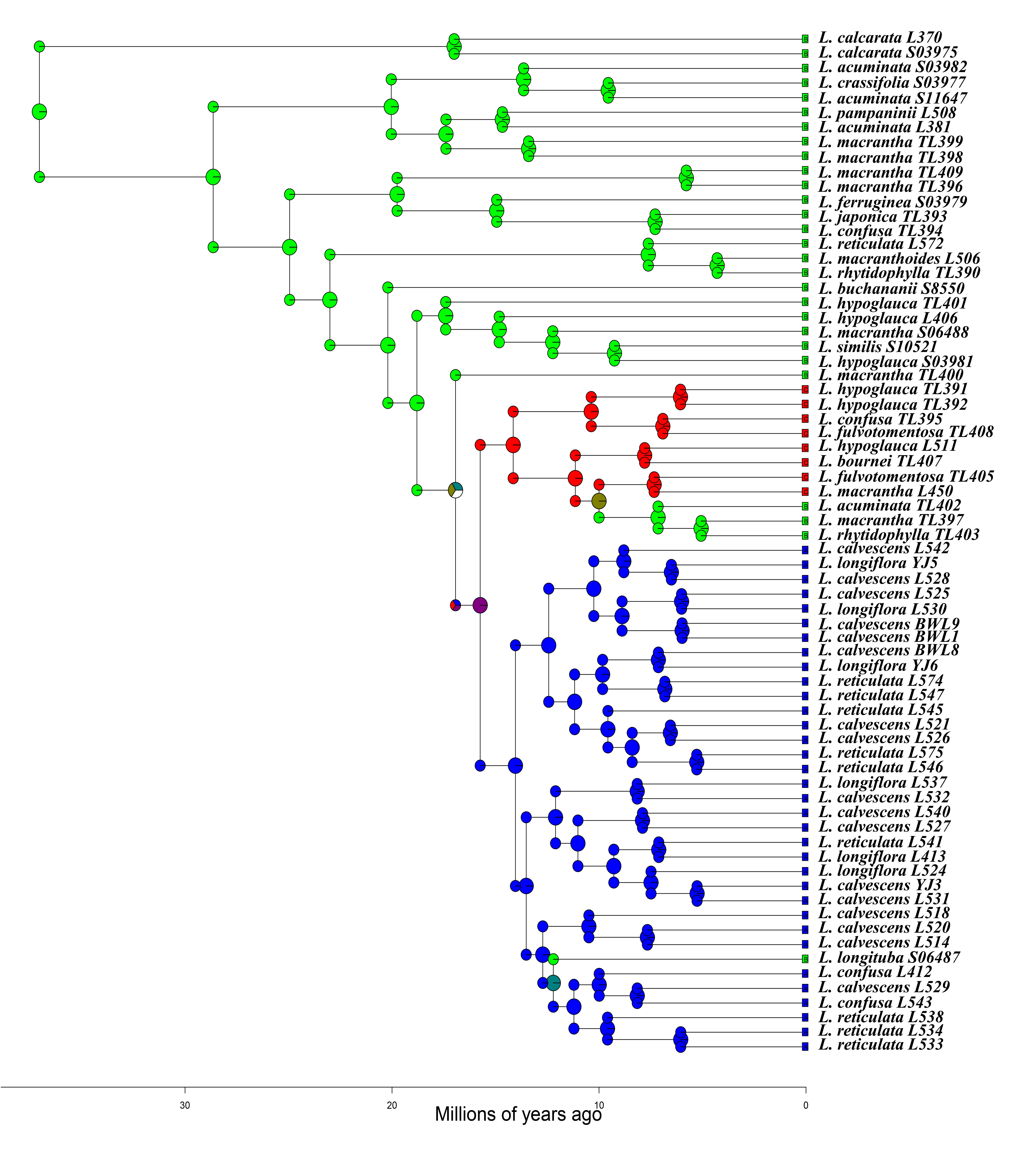

### Supplemental Figure S5

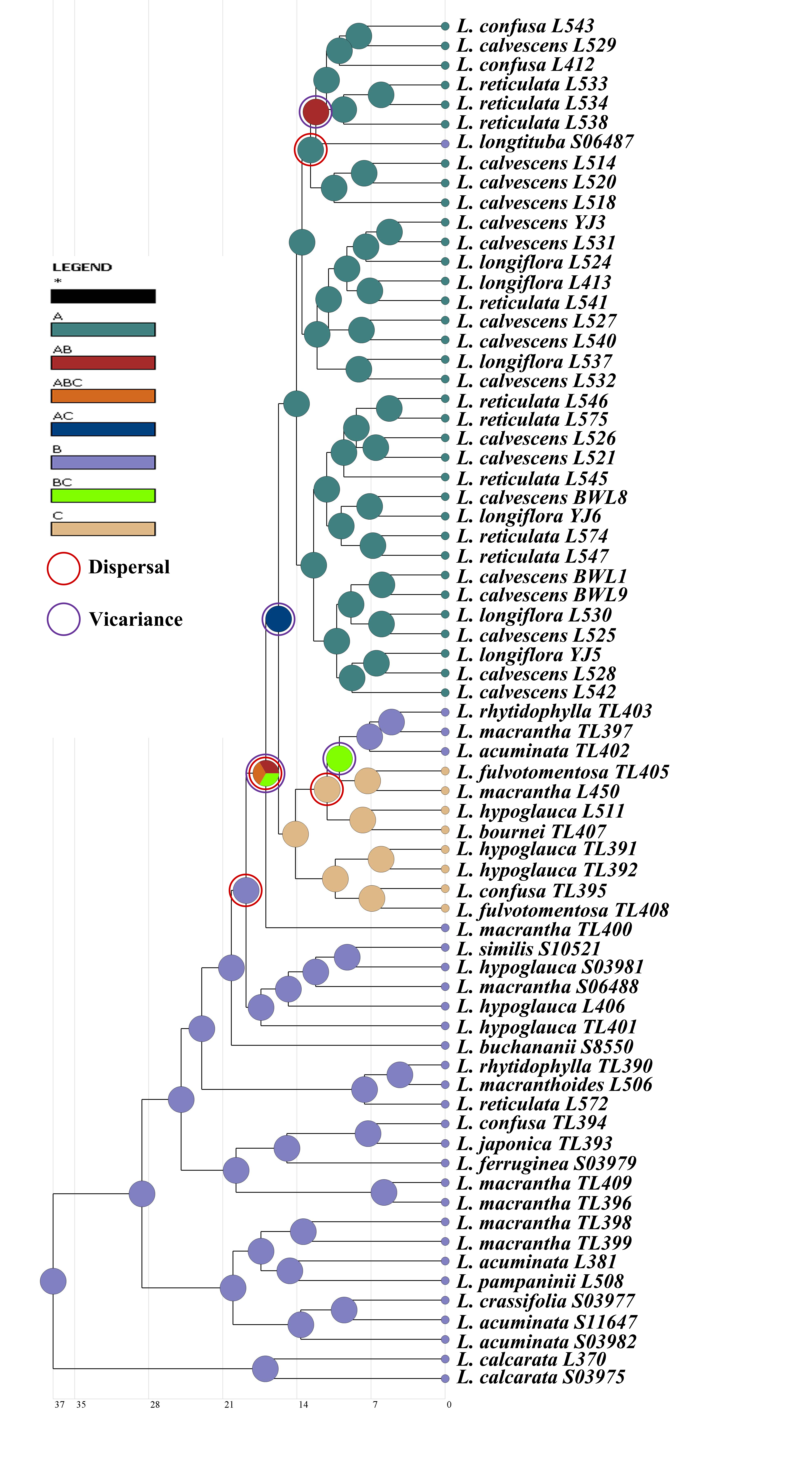

### Supplemental Figure S6

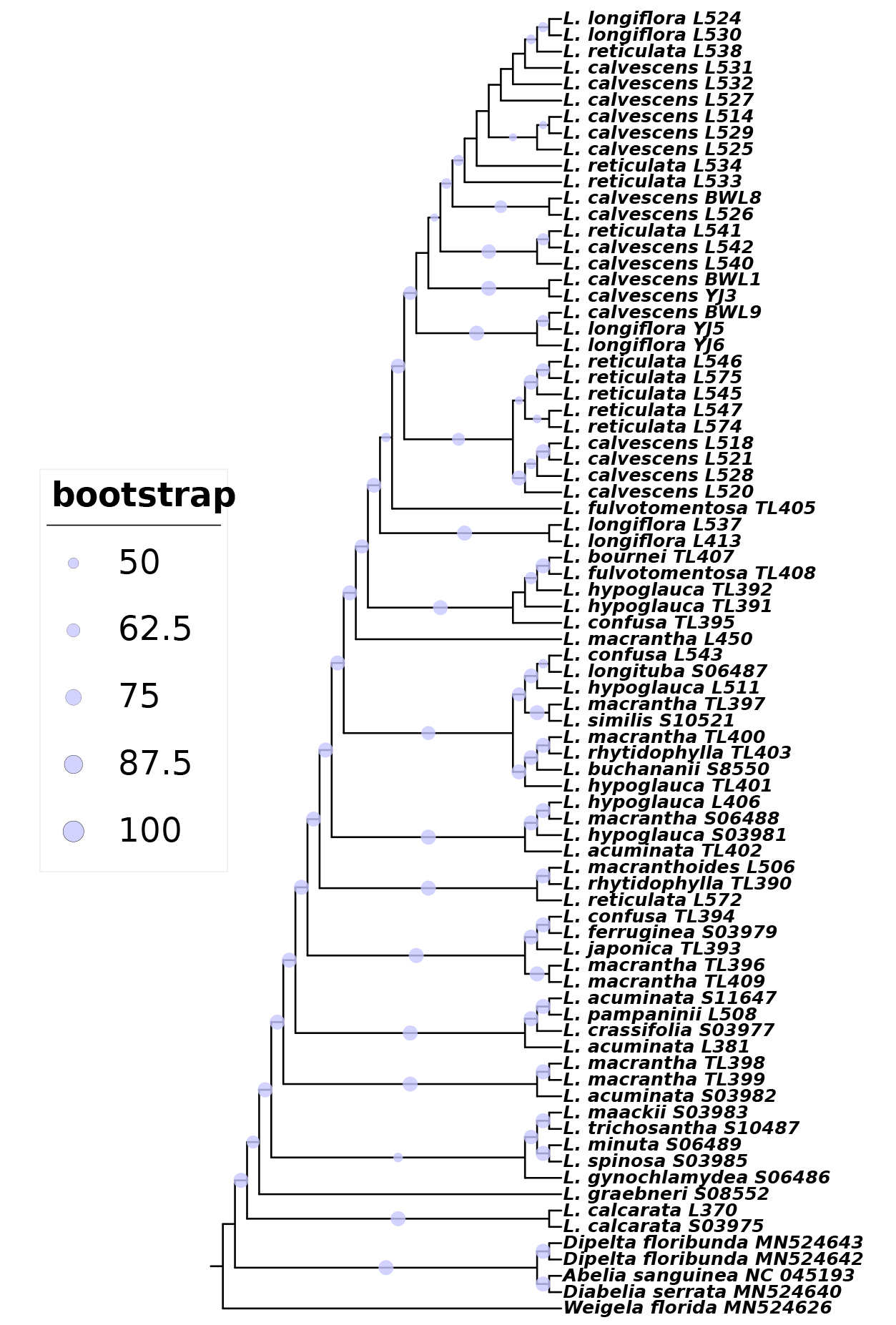

### Supplemental Figure S7

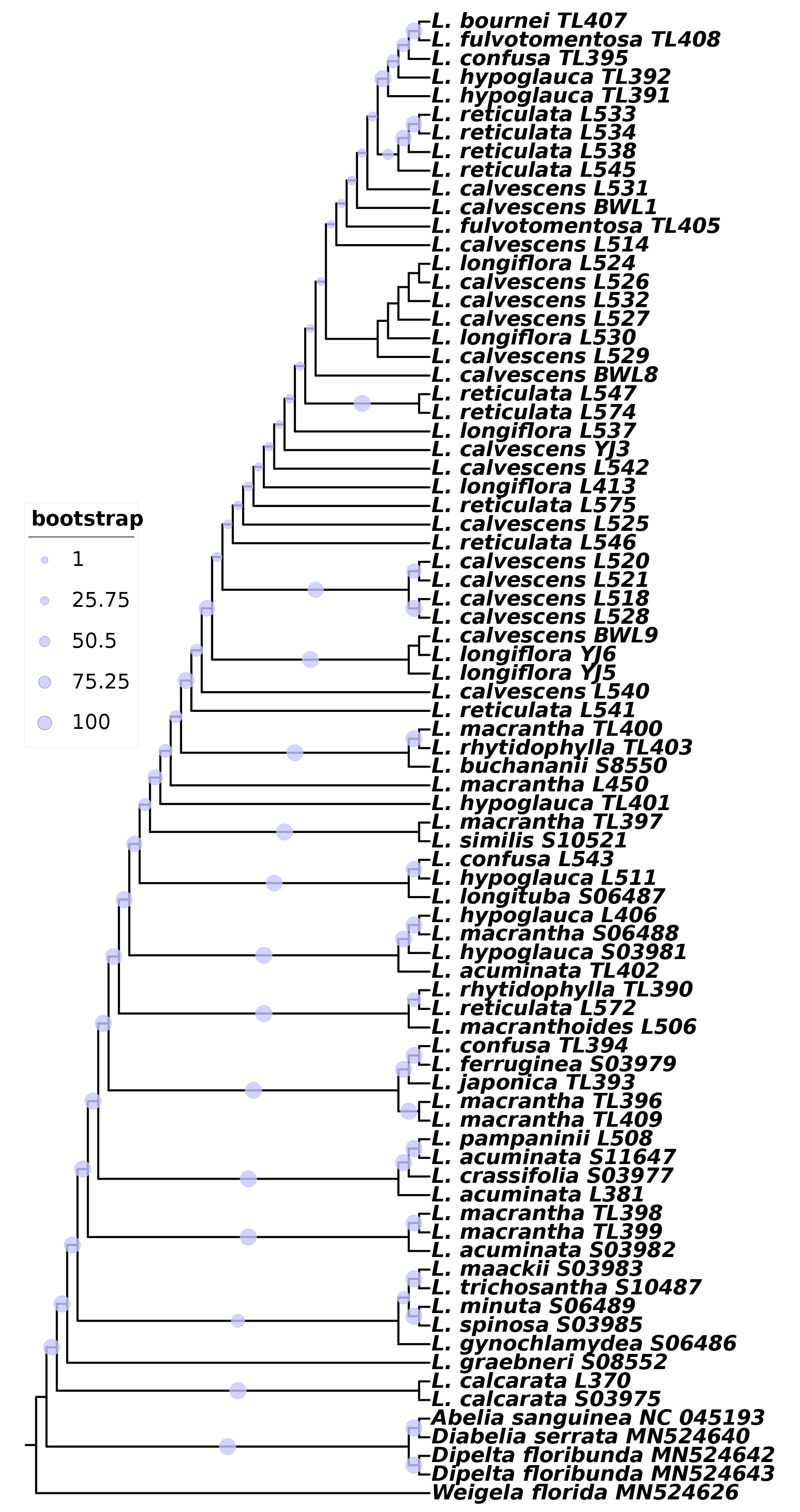

### Supplemental Figure S8

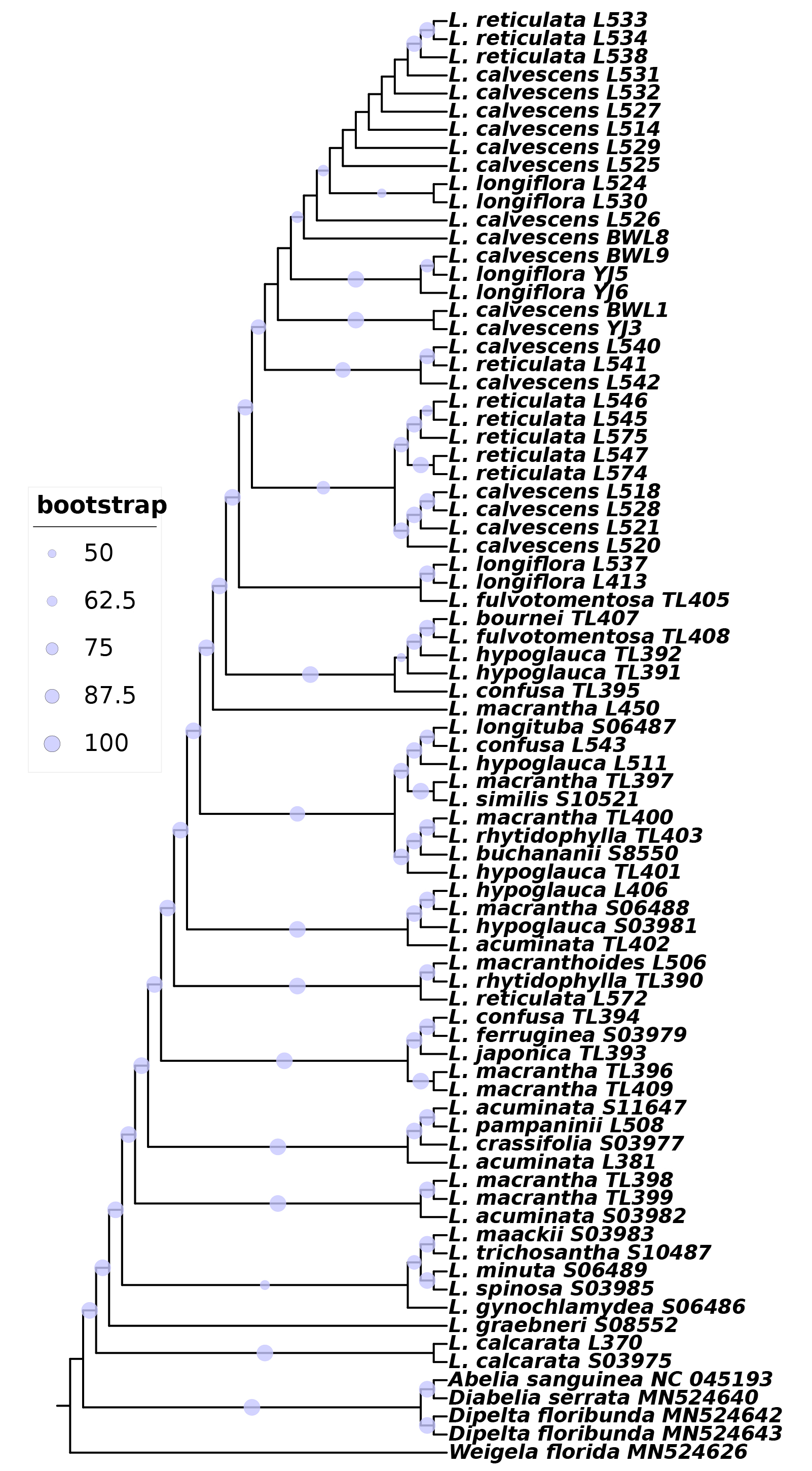

### Supplemental Figure S9

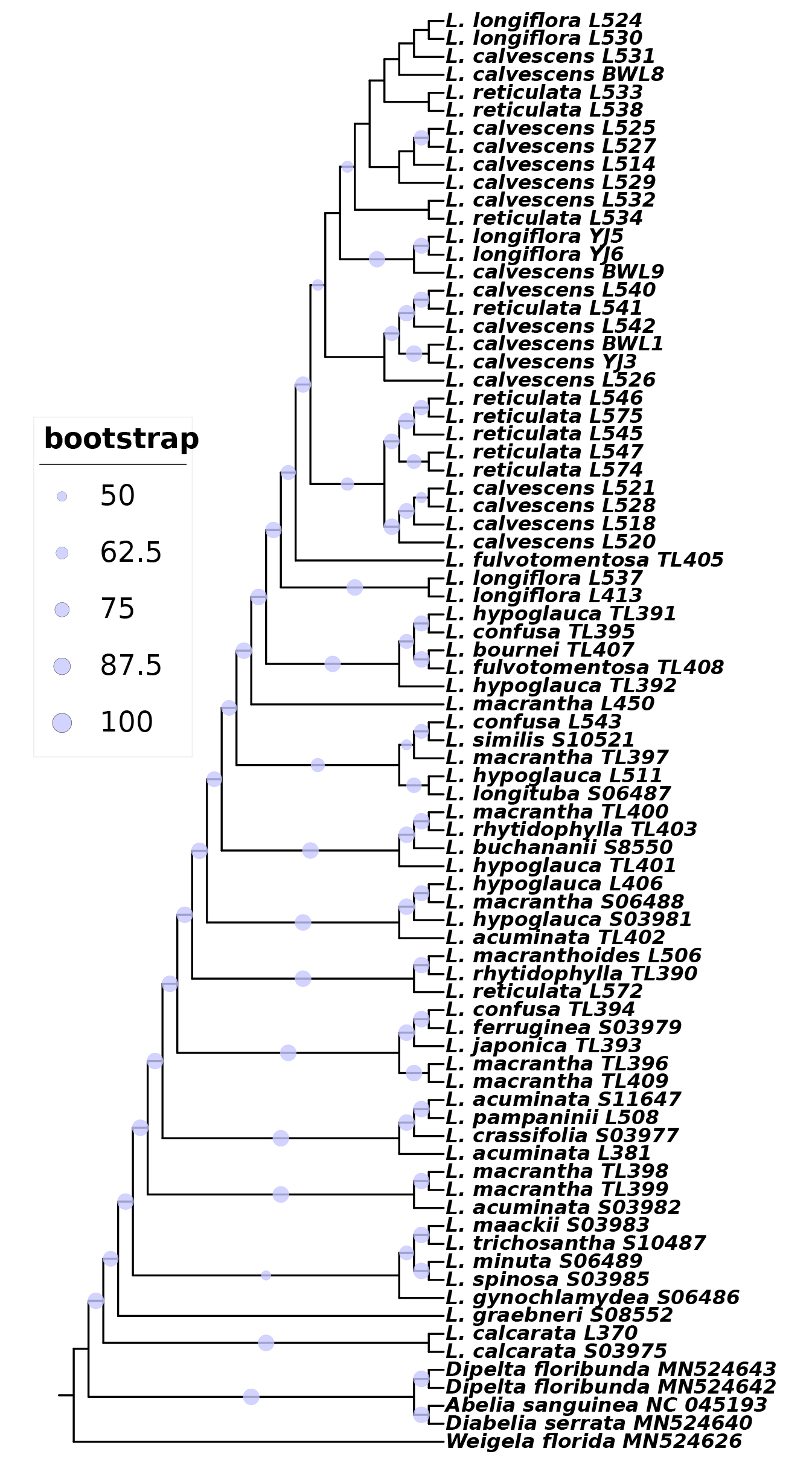

### Supplemental Figure S10

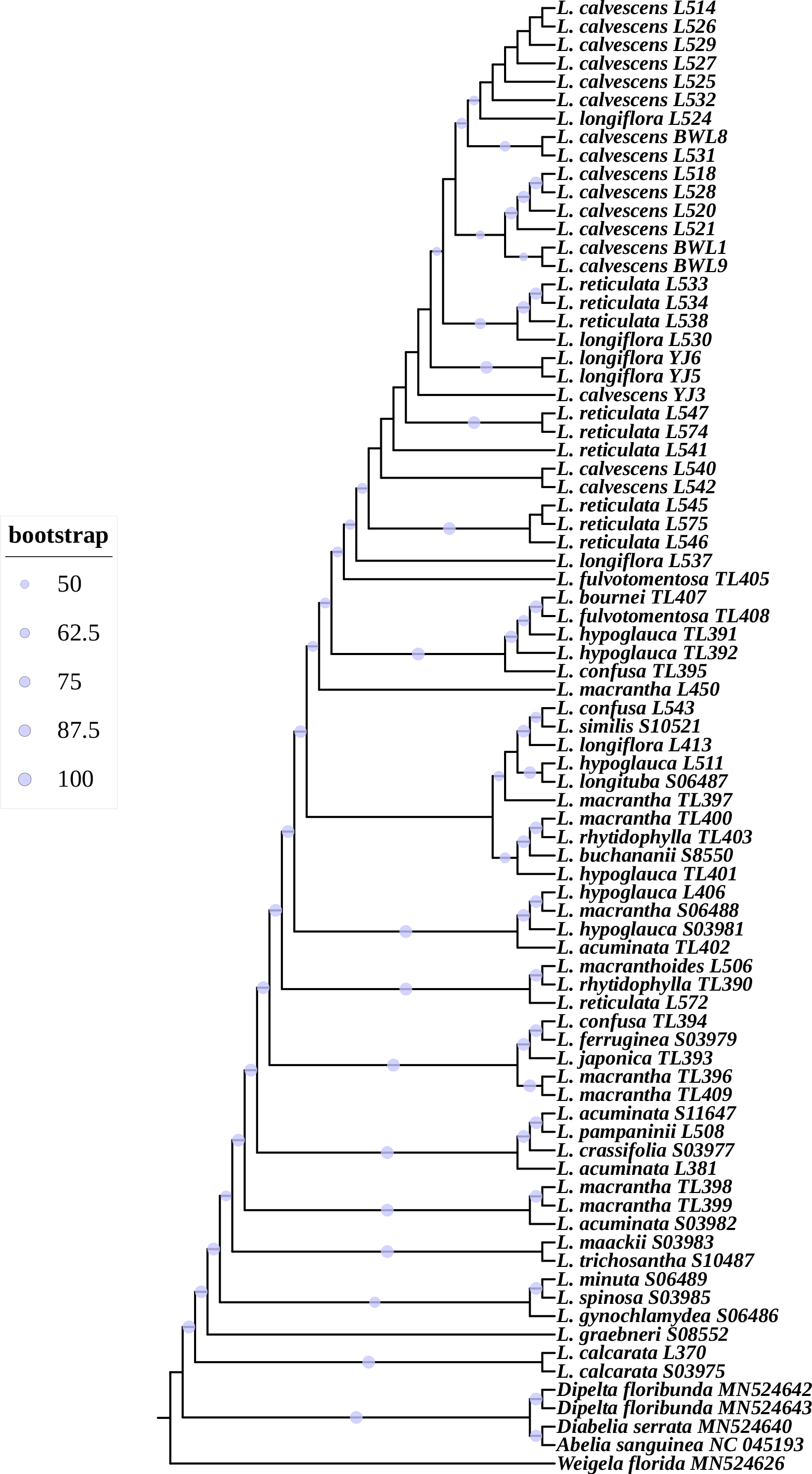

### Supplemental Figure S11

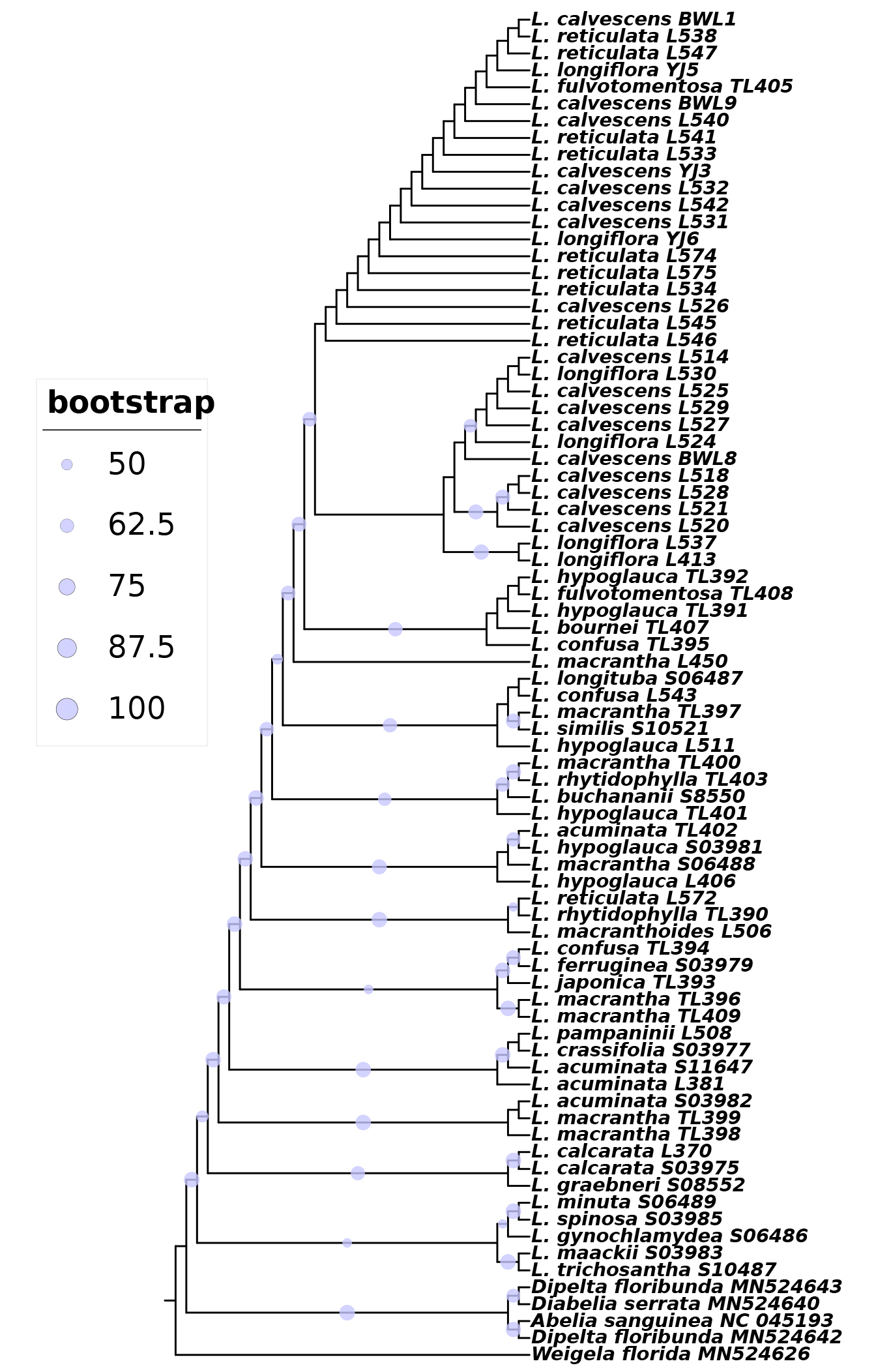
